# Persistent dopamine-dependent remodeling of the neural transcriptome in response to pregnancy and postpartum

**DOI:** 10.1101/2025.02.20.639313

**Authors:** Jennifer C Chan, Giuseppina Di Salvo, Ashley M. Cunningham, Sohini Dutta, Elizabeth Brindley, Benjamin H. Weekley, Ethan Wan, Cindy Zhang, Naguib Mechawar, Gustavo Turecki, Ian Maze

## Abstract

Pregnancy and postpartum experiences represent transformative physiological states that impose lasting demands on the maternal body and brain, resulting in lifelong neural adaptations^1–6^. However, the precise molecular mechanisms driving these persistent alterations remain poorly understood. Here, we used brain-wide transcriptomic profiling to define the molecular landscape of parity-induced neural plasticity, identifying the dorsal hippocampal formation (dHF) as a key site of transcriptional remodeling. Combining single-cell RNA sequencing with a maternal-pup separation paradigm, we additionally demonstrated that chronic postpartum stress significantly disrupts dHF adaptations by altering dopamine dynamics, leading to changes in the dopamine-dependent histone post-translational modification – H3 dopaminylation, with further alterations in transcription, cellular plasticity, and behavior. In human dorsal subiculum, a brain structure within the dHF, we uncovered conserved patterns of parity-dependent alterations in H3 dopaminylation and transcription. We further established the sufficiency of dopamine modulation in regulating these parity-induced adaptations via chemogenetic suppression of dopamine release into the dHF, which recapitulated key epigenomic and behavioral features of parity in virgin female mice. In sum, our findings establish dopamine as a central regulator of parity-induced neuroadaptations in humans and mice, revealing a fundamental transcriptional mechanism by which female reproductive experiences remodel the brain to sustain long-term behavioral adaptations.

## INTRODUCTION

Matrescence – the physical, emotional, hormonal, and social transition to motherhood – is a period of profound transformation that reshapes the maternal body and brain to support pregnancy, birth, and offspring care. While extensive research has focused on acute maternal brain processes that are essential for the onset and maintenance of parenting, precisely how brain adaptations are sustained beyond the postpartum period to persistently influence behavior remains unclear. Recent human neuroimaging studies have revealed that pregnancy experience induces long-lasting alterations in brain connectivity and structure, persisting for years or even decades following birth^1–6^. Animal models examining parity – the state of previously carrying a pregnancy to term – similarly have demonstrated persistent alterations in synaptic remodeling, cell proliferation, and behavioral outcomes^7–14^. However, the specific mechanisms within the extensive repertoire of maternal physiological changes that drive these long-lasting brain adaptations remain poorly understood.

Here, we employed brain-wide transcriptional profiling to systematically characterize the persistent impact of parity on the maternal brain in mice. These analyses surprisingly revealed the dorsal hippocampual formation (dHF) as a key site of sustained plasticity. We then leveraged single-cell RNA sequencing, Cleavage Under Targets and Release Using Nuclease followed by sequencing (CUT&RUN-seq), a postpartum stress paradigm, behavioral assays, chemogenetic approaches, and human brain tissue analyses to identify and validate a dopamine-dependent neuromodulatory process that orchestrates these sustained maternal adaptations. Importantly, these brain-wide datasets will also serve as a critical resource for future investigations into the reproductive state-dependent adaptations that contribute to matrescence.

## RESULTS

### Parity promotes long-lasting transcriptional and behavioral adaptations in maternal brain

To first investigate the persistent effects of female reproductive experiences in maternal brain, we established a timeline to compare primiparous dams (PP), which experienced breeding, pregnancy, parturition, lactation, and pup interactions, to age-matched virgins that lacked reproductive experiences (nulliparous, NP; **Fig. 1a**). Given that reproductive hormones return to baseline levels ∼7-weeks post-conception^15^, we conducted brain-wide transcriptomic profiling 4-weeks following cessation of pup rearing (i.e. 49 days postpartum; dpp), as this time point would be predicted to reflect sustained changes. Following differential expression analyses of 11 brain regions selected based on prior evidence supporting their involvement in maternal behaviors^16,17^ (**Extended Data 1a**), our data revealed a wide-range in the number of differentially expressed genes (DEGs, adj. *p* < 0.05) across brain regions (**Fig. 1b, Extended Data Table 1**). These differences indicated varying levels of transcriptomic sensitivity to parity brain-wide, with few changes observed in the medial prefrontal cortex (mPFC), for example, as compared to the dHF, the region found to be most robustly regulated in our analyses.

**Figure 1.**
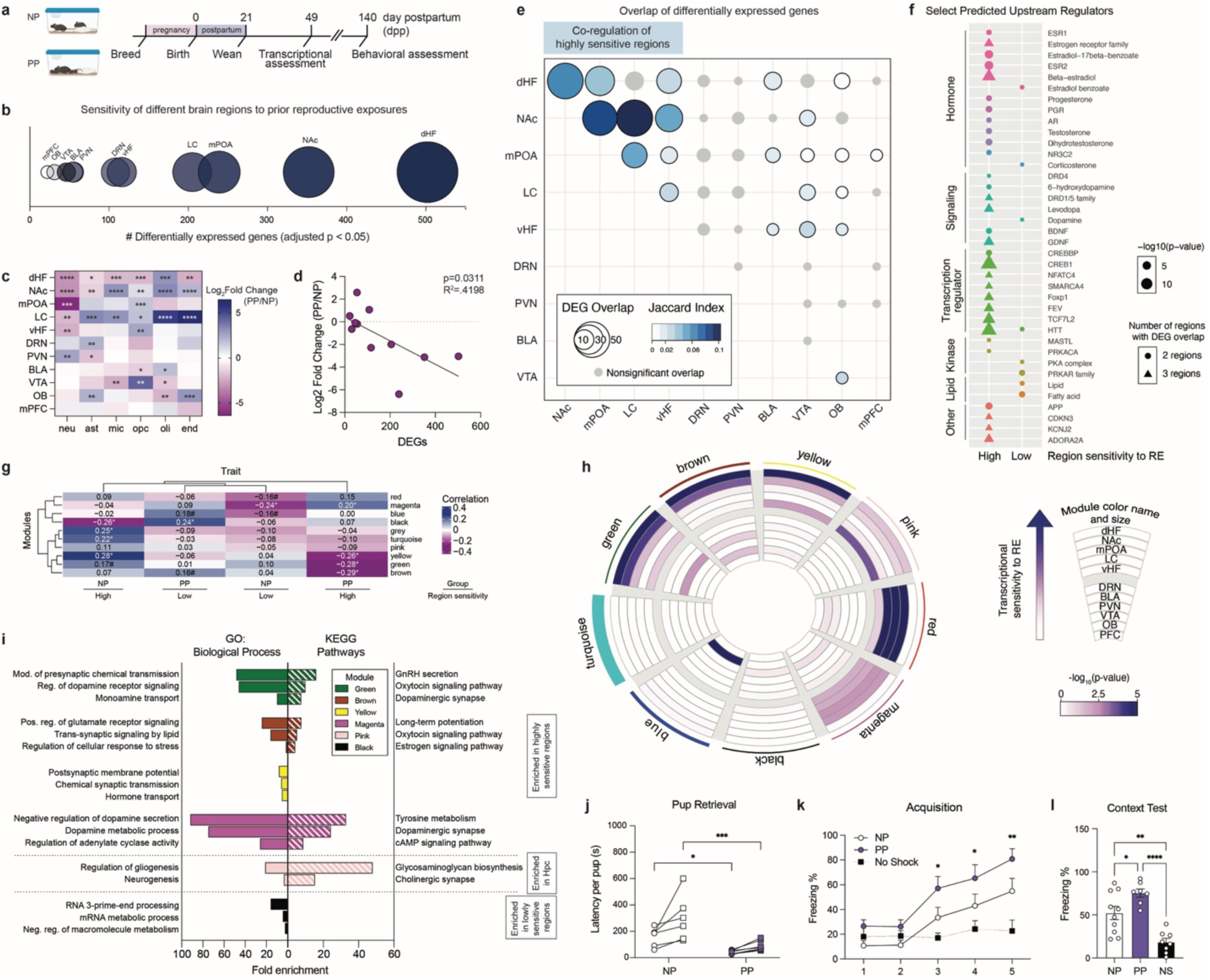
Pregnancy and postpartum states promote lasting transcriptional and behavioral adaptations in the maternal brain. **a)** Experimental timeline comparing primiparous (PP) *vs.* age-matched nulliparous (NP) female mice 1-month after offspring weaning (day postpartum, 49 dpp), and behavioral outcomes at 140 dpp. **b)** Differentially expressed genes (DEGs) per brain region. Bubble size represents total number of NP *vs.* PP DEGs (adj. p < 0.05). **c)** Cell-type deconvolution of bulk RNA-seq data (NP *vs.* PP normalized surrogate proportion variables, Student’s t-test with multiple comparisons correction; *FDR < 0.05, **FDR < 0.01, ***FDR < 0.001, ****FDR < 0.0001). **d)** Downregulation of neuron proportion significantly correlated with number of DEGs per brain region (R^2^ = 0.4198, p = 0.0311). **e)** Overlap of DEGs across brain regions (p < 0.05; gray indicates nonsignificant overlap). **f)** Select predicted upstream regulators of overlapping DEGs across regions with high (dHF, NAc, mPOA, LC, vHF) *vs.* low (DRN, PVN, BLA, VTA, OB, mPFC) sensitivity to parity. **g)** Trait heatmap correlating gene co-expression modules (arbitrary color names, identified by weighted gene correlation network analysis) with group x brain region classification (*p < 0.05, #p < 0.1). **h)** Circos plot for gene co-expression modules, with the size and arbitrary color name of each module indicated by the arc thickness along the perimeter. Enrichment for DEGs per brain region for each module is indicated by the inner rings, with color indicating significance of overlap. **i)** Select GO Biological Processes and KEGG Pathways significantly enriched from gene co-expression modules (FDR < 0.05). **j)** PP dams retrieved both pups significantly faster compared to NP females at 140 dpp (two-way rmANOVA, main effect of group (F(1,11) = 14.47, p = 0.0029), main effect of pup number (F(1,11) = 11.36, p = 0.0062)), with significantly reduced latency for retrieval of pup 1 (Holm-Šídák’s multiple comparisons test, t(22) = 2.301, *adj. p = 0.0312) and pup 2 (Holm-Šídák’s multiple comparisons test, t(22) = 4.098, ***adj. p = 0.0009). N = 6-7 animals/group. **k)** PP dams exhibited increased freezing behavior during the acquisition phase of contextual fear conditioning (two-way rmANOVA, main effect of group (F(2,21) = 4.259, p = 0.028), main effect of preshock (F(2.827, 59.36) = 26.59, p < 0.0001), interaction (F(8,84) = 5.091, p < 0.0001)), compared to no shock controls (Dunnett’s multiple comparisons test, *p ≤ 0.05, **p < 0.01). **l)** During the context recall test, PP dams froze significantly more compared to NP (one-way ANOVA, F(2,22) = 18.72, p < 0.0001; Tukey’s multiple comparisons test, t(22) = 3.640, *p = 0.044), with both groups demonstrating contextual learning compared to no shock controls (PP vs. NS, t(22) = 8.541, ****p < 0.0001; NP vs. NS, t(22) = 5.538, **p = 0.0021). N = 7-10 animals/group. Error bars represent mean ± SEM.

To next assess whether these differential expression patterns could be attributed to changes in shared cell-types, we performed cell-type deconvolution using BRETIGEA, which compares whole transcriptome data to validated brain cell type-specific marker gene sets^18^. We observed changes in cell marker expression for neurons, astrocytes, microglia, oligodendrocyte precursor cells (OPCs), oligodendrocytes, and endothelial cells across brain regions, regardless of the number of DEGs (**Fig. 1c**). These data suggested that cellular changes may occur across brain regions independently of the extent of transcriptomic alterations, potentially reflecting shifts in the proportion of neurons to non-neuronal/glial cells, modifications in synaptic plasticity, or changes in cell identity in response to prior reproductive experiences. Notably, the expression of neuronal markers was found to be downregulated in brain regions displaying the most DEGs (**Fig. 1c**). Furthermore, a significant inverse correlation between reduced neuronal marker expression and the number of DEGs across all brain regions implicated changes in neuronal populations that may underlie the observed transcriptional responses (**Fig. 1d**); note that this relationship was not observed for other cell-types (**Extended Data 1b-f**). These findings nicely align with human neuroimaging studies, which have reported reduced gray matter in mothers even years after birth, a phenomenon that has been proposed to refine maternal neural circuits to optimize caregiving behavior^1,2,4,19^.

Based on these findings, we hypothesized that such shared neuronal marker alterations may indicate common upstream mechanisms driving regional-specific transcriptional responses. To test this, we first compared DEGs across brain regions to identify common gene sets that were altered by parity status, which led to the observation that the greatest extent of overlap occurs in those regions displaying the highest levels of transcriptional sensitivity (**Fig. 1e**). Based on these analyses, we then grouped the 11 brain regions according to the number of DEGs, the degree of overlap with other regions, and significant fold changes in neuronal marker expression, based on the premise that these regions may be co-regulated to elicit convergent transcriptional responses to parity (referred to as “high sensitivity” regions: dHF, nucleus accumbens (NAc), medial preoptic area (mPOA), locus coeruleus (LC), ventral hippocampus (vHF)). Brain regions lacking these criteria were classified as “low sensitivity” regions (dorsal raphe nucleus (DRN), paraventricular nucleus of the hypothalamus (PVN), basolateral amygdala (BLA), ventral tegmental area (VTA), olfactory bulb (OB), mPFC). Next, we explored the predicted upstream regulators of overlapping DEGs in “High” versus “Low” sensitivity brain regions, which revealed several upstream regulators that were grouped into common molecular classes, including hormones, neurotransmitter/neurotrophin signaling molecules, transcriptional regulators, and others (**Fig. 1f**). While both High and Low sensitivity regions were found to share upstream regulators within these molecular classes, High sensitivity regions displayed greater enrichment for estrogen, progesterone, testosterone, dopamine, and other ligands with similar receptor affinity or structural homology. In contrast, regulators that were classified within the lipid category were uniquely associated with Low sensitivity regions.

To further explore the regulatory networks that distinguish High versus Low sensitivity regions, we performed weighted gene co-expression network analysis (WGCNA) across all 11 brain regions^20^, resulting in nine co-expression modules (**Fig. 1g-j**). Module-trait correlations, in which the trait refers to a combination of group (NP *vs.* PP) and regional sensitivity (High *vs.* Low), identified significant correlations between High sensitivity regions in PP females and the brown, green, yellow, and magenta modules (**Fig. 1g**; henceforth labeled ‘parity-sensitive modules’). We then examined enrichment of DEGs within each module, with an extended focus on the parity-sensitive modules. Consistent with our module-trait correlational analysis, parity-sensitive modules displayed significant enrichment for DEGs from dHF (**Fig. 1h**; Fisher’s exact tests: brown, p < 10^-16^; green, yellow, p < 10^-6^), NAc (brown, green, p < 10^-3^; magenta, p < 0.01; yellow, p < 0.05), mPOA (green, p < 0.05; magenta, p < 0.01), LC (magenta, p < 0.05), and vHpc (magenta, p < 0.05; yellow, p < 0.01). Functional annotation analysis of the genes from these modules highlighted their significant enrichment in signaling pathways related to neuromodulators, such as estrogen (yellow, brown) and dopamine (green, magenta), in agreement with our upstream regulator analyses (**Fig. 1i, Extended Data 1k-l**). In contrast, Low sensitivity regions were significantly associated with the black module, which was enriched for genes associated with RNA and macromolecule processing pathways (**Fig. 1g**, **1i**).

Building on these transcriptomic findings, we next investigated whether parity leads to sustained functional alterations in PP females. To evaluate the extent of such persistence, we performed behavioral assessments at 140 dpp, ∼16-weeks after pup weaning. Behavioral tasks were selected to specifically assess functions associated with brain regions displaying the most robust transcriptional changes, including maternal behavior, learning, and memory. To assess maternal behavior, we tested pup retrieval in the home cage by placing two donor pups in opposite corners away from the nest. PP dams were observed to retrieve pups with significantly reduced latency as compared to NP females (**Fig. 1j**), which is consistent with previous findings, though at a more extended timepoint in our studies^21^. Next, to investigate the functional impact of dHF transcriptional changes, we utilized a contextual fear conditioning paradigm. Female mice were habituated to the training context before receiving five shocks (2-seconds each, 0.7 mA) with 90-second intertrial intervals. Both NP and PP females demonstrated significant training acquisition, as indicated by increased freezing compared to no-shock controls (**Fig. 1k**). Notably, PP females exhibited enhanced acquisition, with significantly increased freezing observed by the third shock, while NP females only displayed significant freezing at the fifth shock (**Fig. 1k**). During the context test conducted 24-hours later, both groups demonstrated recall, with PP females showing higher freezing responses than NP animals (**Fig. 1l**). Given the involvement of these brain regions in anxiety- and depressive-like behaviors, we also assessed affective phenotypes using the open field, light-dark box, and forced swim tests, however, no significant differences were observed (**Extended Data 2a-h**). These results suggested that the transcriptional changes observed in parity-sensitive brain regions may promote maternal and dHF-related behavioral outcomes, as has been reported previously^7,10,12,14,22–28^. To account for the potential influence of estrous cycling on behavioral changes, vaginal swabs were collected immediately following testing, and behavioral analyses were stratified by estrous stage. No significant effects of estrous stage were observed (**Extended Data 2i-n**), supporting parity status as the primary driver of the observed behavioral differences.

### Pregnancy and postpartum experiences promote persistent maternal neuroplasticity

Since our paradigm comparing NP *vs.* PP females could not distinguish between the discrete reproductive event(s) that may be responsible for the sustained transcriptional alterations observed, we next sought to isolate those key events throughout the female reproductive window. To do so, we compared females that were successfully bred to males but did not become pregnant (Mating Only), females that experienced pregnancy and parturition but did not transition to postpartum due to pup removal on the day of birth (Pregnancy Only), and virgin females exposed to 21-days of donor pup interactions (Pup Sensitized). These groups were analyzed in parallel with age-matched NP and PP females to delineate the contributions of specific reproductive and maternal experiences (**Fig. 2a**). Consistent with previous studies^29^, four days of pup exposures sensitized virgin females to maternal behavior, as evidenced by increased crouching over pups and decreased pup avoidance by day 4 (**Extended Data 3a**). To determine the processes conferring sustained transcriptional sensitivity, we focused our investigations on the dHF due to its pronounced gene expression response to parity. When comparing NP *vs.* PP gene expression profiles, we observed that the Pregnancy Only group most closely resembled that of the expression profile of PP females, suggesting that pregnancy is a primary driver of dHF neuroplastic changes (**Fig. 2b, Extended Data 3b**). However, the magnitude of fold changes did not match that of the PP group, indicating that additional experiences, likely during postpartum, are also necessary to fully establish parity-induced changes in the dHF.

**Figure 2.**
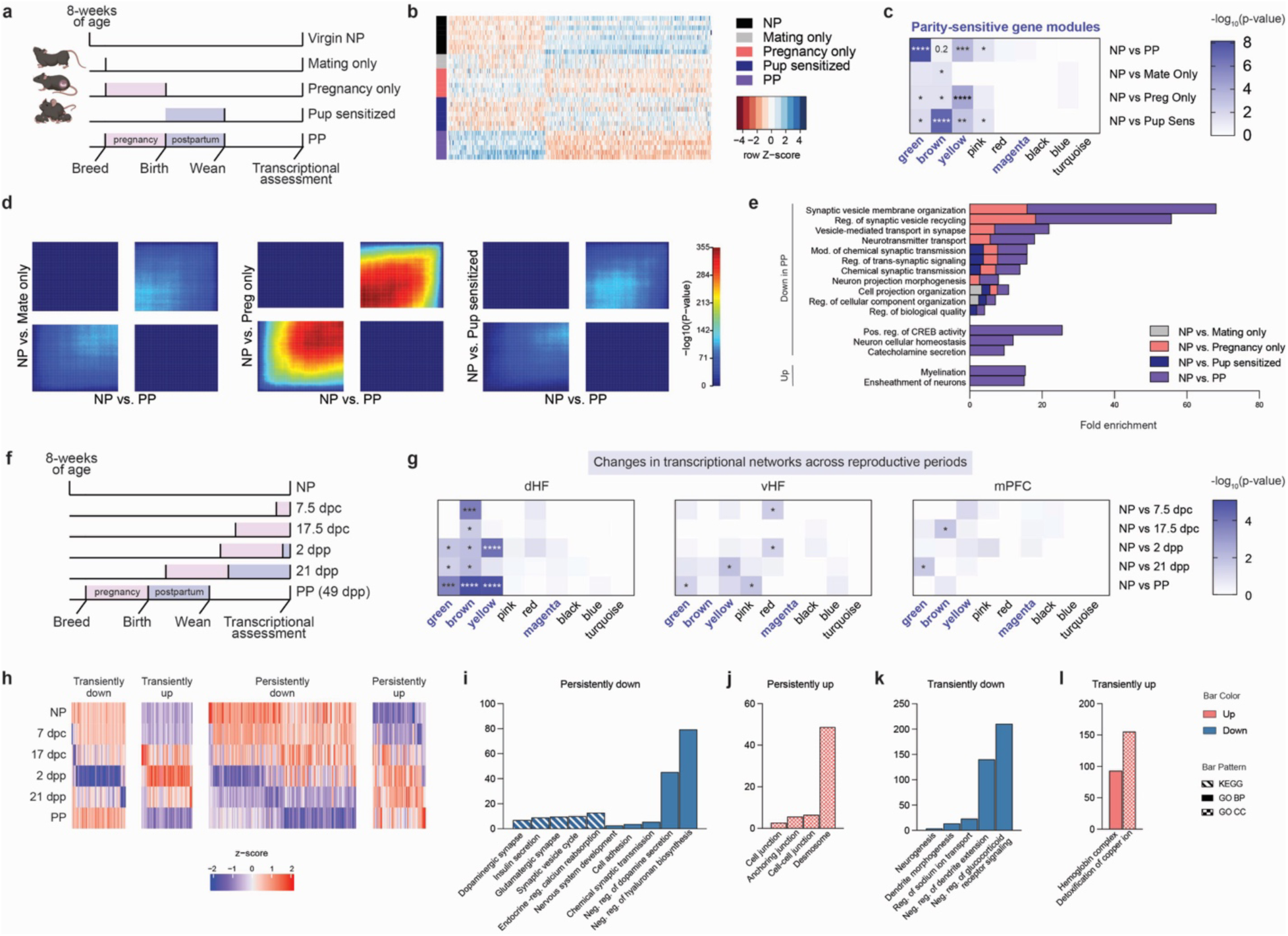
Parity adaptations in the dHF transcriptome integrate pregnancy and postpartum experiences. **a)** Experimental design to examine the contribution of discrete female reproductive events. **b)** Heatmap of differential expression profiles of top 500 genes, by ascending p-values, between NP *vs.* PP (N=3-7/group). **c)** Overlap of DEGs (*vs.* NP, adj. p < 0.05) for each gene co-expression module, with color representing extent of significance (*p < 0.05, **p < 0.01, ***p < 0.001, ****p < 0.0001). High parity sensitivity modules are indicated in bold. **d)** Threshold-free comparison of indicated comparisons by rank-rank hypergeometric overlap. Pixels represent the overlap of differential expression profiles indicated, with color representing extent of significance. The lower left and upper right quadrants represent concordant gene regulation. **e)** Shared changes in biological processes (identified through GO term analysis) observed following individual reproductive exposures with PP, along with processes exclusive to the combined experiences in PP (FDR < 0.05). **f)** Experimental design to examine the trajectory of PP dHF gene expression changes across pregnancy and postpartum periods (dpc: days post-conception; dpp: days postpartum). **g)** Enrichment of DEGs (*vs.* NP, adj. p < 0.05) with parity-sensitive gene co-expression modules across reproductive time points, examining extent of significant overlap in brain regions exhibiting high (left, dHF), moderate (middle, vHF), and low (right, mPFC) transcriptional sensitivity to parity status (*p < 0.05, **p < 0.01, ***p < 0.001, ****p < 0.0001). N=6/group. **h)** Impulse time course analysis of dHF gene expression identified patterns of transient *vs.* persistent up- and downregulation across pregnancy and postpartum (FDR < 0.05). **i-l)** Select GO terms and KEGG pathways significantly enriched for dHF genes exhibiting patterns of **(i)** persistent downregulation, **(j)** persistent upregulation, **(k)** transient downregulation, and **(l)** transient upregulation across reproductive stages (FDR < 0.05).

While visualization of differential expression profiles in the Mating Only and Pup Sensitized groups did not mimic PP changes, comparisons to NP did identify significant DEGs across all groups, with the most substantial changes observed in the Pup Sensitized group, followed by the Pregnancy Only, and Mating Only groups (**Extended Data 3c-e, Extended Data Table 2**). Overlapping DEGs induced by discrete reproductive events significantly intersected with DEGs in PP animals, suggesting that while pregnancy is the primary driver of parity-induced effects, each reproductive event contributes, in part, to this process (**Extended Data 3f**). Next, we overlapped DEGs from each comparison with genes from parity-sensitive modules (from **Fig. 1**). Significant overlap was observed across all groups, with Pregnancy Only and Pup Sensitized groups displaying the strongest enrichment, indicating that these experiences play a key role in shaping persistent dHF alterations (**Fig. 2c**). Thus, despite the lack of similarity between the overall expression profiles between Pup Sensitized *vs.* PP, gene expression changes induced by pup interactions likely converge on similar regulatory networks, albeit through different DEGs, which are critical for parity-driven transcriptional alterations. To further explore these relationships, we performed threshold-free Rank-Rank Hypergeometric Overlap (RRHO) analyses^30^, which demonstrated transcriptional concordance across experiences, with the strongest overlap observed in pregnancy (**Fig. 2d**). These findings suggest that reproductive exposures converge, or act additively, to drive the full extent of brain transcriptional changes. Furthermore, comparing GO term analyses of DEGs for each group revealed significant enrichment in overlapping biological processes, with many related to synaptic signaling and plasticity pathways (**Fig. 2e**). Notably, certain pathways were unique to the combination of reproductive exposures, including changes in CREB activity, catecholamine secretion, and myelination, indicating that these regulatory processes depend on multiple reproductive events to achieve full induction (**Fig. 2e**).

Given that postpartum experiences enhanced changes that were initiated during pregnancy, we conducted a controlled time course study to examine these transcriptional dynamics across gestational and postpartum periods in age-matched females (**Fig. 2f**). Pregnancy timepoints were selected to represent the anabolic and catabolic phases of gestation, corresponding to early (7.5 days post-conception, dpc) and late (17.5 dpc) stages, respectively. Postpartum timepoints were chosen to capture the immediate effects of hormonal withdrawal following parturition (2 dpp), as well as a later stage prior to pup weaning (21 dpp). Based on the hypothesis that parity-induced transcriptional alterations are progressively enhanced across reproductive stages, we first compared DEGs at each timepoint (*vs.* NP) to genes from parity-sensitive modules identified previously. DEGs from all comparisons significantly overlapped with the brown module (**Fig. 2g, left; Extended Data Table 3**), suggesting that pathways enriched in this module are central to dHF programming. Enrichment in the green and yellow modules was predominantly found to be induced by postpartum exposures, potentially amplifying the effects of the pathways driven by the brown module (**Fig. 2g, left**). To assess the regional specificity of these dynamic changes, we analyzed bulk RNA-seq data from the vHpc, a region that displayed moderate sensitivity to parity status (**Fig. 2g, middle; Extended Data Table 4**), along with the mPFC, which exhibited few transcriptional alterations (**Fig. 2g, right; Extended Data Table 5**). These regions exhibited limited overlap of DEGs with parity-sensitive module genes, further emphasizing the specificity of the pathways that drive long-term adaptations in brain regions displaying heightened sensitivity to parity.

To then characterize cumulative transcriptional changes that occur across reproductive stages, we performed a time course analysis using the ImpulseDE2 package to detect genes displaying sustained or transient expression alterations^31^. This analysis revealed a prominent set of genes that displayed persistent and progressive downregulation, with more pronounced alterations during postpartum (**Fig. 2h**). These genes were significantly enriched for processes related to glutamatergic synapses, dopaminergic signaling, and endocrine-related pathways (**Fig. 2i**). Conversely, genes with persistent upregulation were found to be enriched for pathways associated with cell junction dynamics (**Fig. 2j**), which is consistent with overall prolonged changes in synaptic plasticity and transmission. Additionally, we identified gene sets displaying transient alterations, including downregulation of neurogenesis-related pathways, and upregulation of copper ion metabolism (**Fig. 2k-l**); these data are consistent with other studies reviewed here^32,33^. In total, these findings highlight postpartum as a key window for reinforcing parity-induced alterations.

### Chronic maternal stress during postpartum disrupts dHF adaptations

We next examined whether postpartum perturbations may disrupt pathways that reinforce parity-induced alterations in dHF. As becoming a new parent represents a significant source of stress, we implemented a maternal stress paradigm that robustly increases stress hormone levels and disrupts key postpartum experiences, including nursing and pup interactions, from 10–20 dpp (**Fig. 3a, Extended Data 3g**). During this period, dams were separated from their pups for 3-hours daily and were provided with limited nesting materials until pup weaning (Stress PP). As litter size represents a potential confounding factor, the time period selected reduces the impact of maternal stress on pup mortality that can occur immediately following birth. By 10 dpp, pups were more durable, as reflected by the absence of litter size differences between Control and Stress dams (**Extended Data 3h**). Accordingly, no differences in postpartum weights were observed, suggesting that this postpartum stress paradigm does not induce major metabolic alterations in the dams (**Extended Data 3i**). To assess the extent of parity adaptations in Stress PP, we compared this group to NP and Control PP animals. Principal components analysis (PCA) of dHF transcriptional profiles revealed distinct clustering of NP and Control PP groups (reproducing our prior analyses), while Stress PP samples were found to cluster intermediately between the two groups (**Fig. 3b**). Similarly, hierarchical clustering of all DEGs resulted in intermingling of NP and Stress PP samples (**Fig. 3c, Extended Data Table 6**), suggesting that postpartum stress disrupts the extent of parity alterations in dHF. To determine whether genes induced by parity are reversed by stress, we then compared DEGs with opposing directionality in NP *vs.* Control PP and Control PP *vs.* Stress PP groups (**Fig. 3d**). Approximately 85% of genes altered by maternal stress overlapped with parity-induced gene expression changes. KEGG pathway analysis of DEGs revealed shared enrichment across several pathways in both comparisons, with maternal stress reversing gene expression associated with long-term potentiation, dopaminergic synapses, oxytocin signaling, and other processes (**Fig. 3e**).

**Figure 3.**
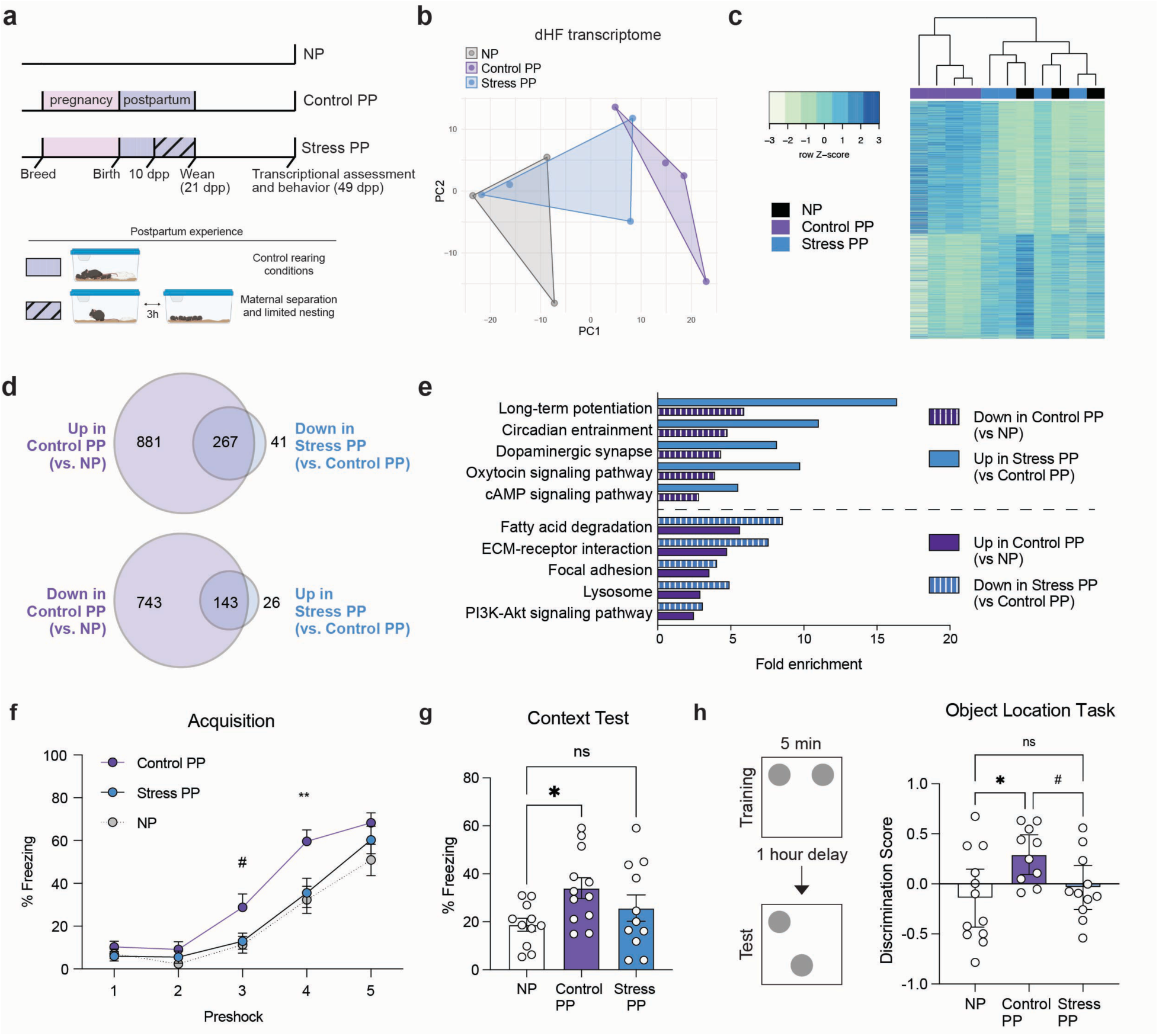
Postpartum stress disrupts long-term maternal dHF adaptations. **a)** Postpartum stress paradigm, wherein dams were provided with limited nesting and subjected to pup separation 3 hours daily between 10-20 dpp, followed by behavioral and transcriptomic assessments at 49 dpp. **b)** Principal components analysis of dHF transcriptomes, and **c)** heatmap of all significant DEGs (adj. p < 0.05) with hierarchical clustering. **d)** Venn Diagrams showing ∼20% DEGs altered in Control PP (*vs.* NP) are significantly disrupted by postpartum stress (adj. p < 0.05). **e)** Parity-dependent enrichment in select KEGG pathways are significantly reversed in Stress PP (FDR < 0.05). **f)** Control PP dams froze more during the acquisition phase of contextual fear conditioning (two-way rmANOVA, main effect of group (F(2,31) = 4.662, p = 0.017), main effect of preshock (F(3.024,93.75) = 129.8, p < 0.0001), interaction (F(8,124) = 2.406, p = 0.091)), compared to NP and Stress PP (Tukey’s multiple comparisons test, #p ≤ 0.1, *p < 0.05, **p < 0.01). **g)** During the context recall test, NP froze significantly less than Control PP (one-way ANOVA, F(2,31) = 3.151, p = 0.05; Tukey’s multiple comparisons test, t(31) = 3.54, *p = 0.0456), but not Stress PP (ns, p > 0.05). N = 11-12 animals/group. **h)** Control PP, but not Stress PP, dams spent more time investigating the moved object in the object location task compared to NP (one way ANOVA, F(2,30) = 4.045, p = 2.0278; Holm-Šídák’s multiple comparisons test: NP vs. Control PP, t(30) = 2.764, *adj. p = 0.0288; Control PP vs. Stress PP, t(30) = 2.046, #adj. p < 0.1) N=10-12 animals/group. Error bars represent mean ± SEM.

Since our pathway analyses indicated changes in long-term potentiation, a form of neuronal plasticity that is important for learning and memory behaviors^34^, we further focused our behavioral assessments on contextual fear conditioning. Consistent with prior results, Control PP females exhibited enhanced acquisition and context recall, as demonstrated by significantly increased freezing behavior compared to NP animals. In contrast, however, Stress PP females showed no significant differences *vs.* NP mice (**Fig. 3f-g**). To further evaluate dHF-dependent function, we next assessed behavior using the object location task. In this test, animals were initially trained by allowing them to explore two identical objects, followed by one-hour removal from the arena. During the test phase, one object was displaced, and the animals’ ability to identify the moved object was assessed. Given prior findings that adult females require 10 minutes of training for reliable discrimination^35^, we first used this duration to confirm that all groups could successfully identify the moved object (**Extended Data 3j**). To next assess whether Control PP females exhibit enhancements in this task, we reduced the training time to 5 minutes, a duration that is sufficient for learning in adult males and preadolescent female mice, but not in adult females^35^. Under these conditions, Control PP females retained the ability to discriminate the novel location, whereas NP and Stress PP females did not, reflecting the same pattern observed in contextual fear conditioning, which was not attributed to estrous stage or locomotion (**Extended Data 3k-l)**.

### Cell-type specific transcriptional alterations reveal dopaminergic modulation underlying maternal dHF plasticity

To elucidate the mechanisms underlying parity-induced dHF plasticity, we next leveraged the transcriptomic and behavioral signatures shared between NP and Stress PP groups. This postpartum stress paradigm resolves key windows during which disruptions to parity adaptations occur, thereby providing greater temporal resolution of the mechanisms required to sustain these alterations. Building on our findings, we next performed single-nuclei (sn) RNA-sequencing on these three groups to identify the dHF cell-types that integrate hormone and neurotransmitter signaling to drive parity-related plasticity. Following rigorous quality control assessments to remove confounding sources of variation, such as mitochondrial mapping percentage, doublets, and cell-type contamination **(Extended Data 4a-f)**, we retained 109,334 nuclei for downstream analysis (NP = 37,070; Control PP = 35,631; Stress PP = 36,633). Cell-type annotation was performed using both manual curation and label transfer from validated hippocampal datasets from the Broad Institute and Allen Brain Atlas^36,37^. This resulted in 16 distinct clusters (**Fig. 4a, Extended Data 4g-i**), including excitatory neuron subtypes (dentate gyrus, CA1, CA3, subiculum, mossy cells; 43.3% of total), GABAergic neurons (GABA.1–GABA.4; 9.1% of total), neural progenitors (neural stem cells, radial glia-like cells; 9.7% of total), and non-neuronal/glial populations (astrocytes, oligodendrocytes, OPCs, microglia, immune cells; 37.9% of total).

**Figure 4.**
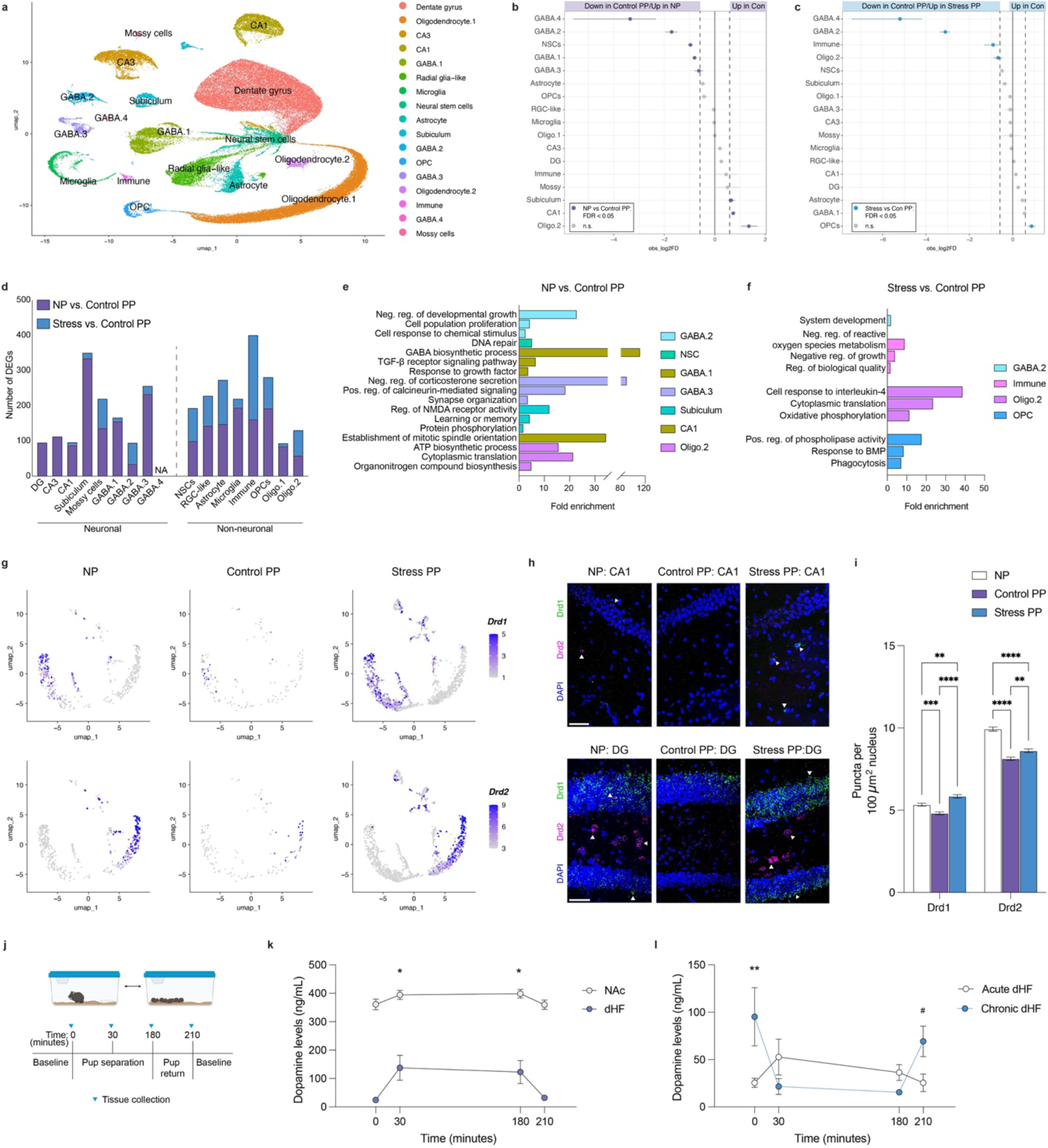
Parity-driven dHF plasticity is suppressed by elevated dopamine signaling. **a)** UMAP representation of cell clusters (109,334 total cells) from dHF tissues, colored by major cell types (N = 6 NP, 4 Control PP, 5 Stress PP). **b, c)** Point range plots depicting significant changes in proportions of major cell-type clusters between **(b)** NP vs. Control PP, and **(c)** Stress PP vs. Control PP (FDR < 0.05 and |log_2_Differerence| > log_2_(1.5)). **d)** Number of DEGs observed for each major cell-type following pseudobulk analysis (p < 0.05 and |log_2_(Fold Change)| > log_2_(1.5)). **e, f)** GO term analyses (Biological Process) for DEGs identified in cell clusters with significant proportional differences between **(e)** NP vs. Control PP, and **(f)** Stress vs. Control PP (FDR < 0.05). **g)** Subclustering of GABA.2 neuronal population revealed reductions in *Drd1* (top) and *Drd2* (bottom) expression in Control PP. **h)** Representative images of RNAScope for *Drd1* and *Drd2* mRNAs in dorsal CA1 (top) and dentate gyrus (DG, bottom). Scale bars, 50 μm. **i)** Quantification of *Drd1* and *Drd2* mRNA puncta in CA1 and DG nuclei (two-way ANOVA, main effect of group (F(2,82142) = 65.81, p < 0.0001), main effect of gene (F(1,82142) = 1771, p < 0.0001), interaction (F(2,82142) = 39.99, p < 0.0001); Tukey’s multiple comparisons test, **p < 0.01, ***p < 0.001, ****p < 0.0001). N = 5 animals/group. **j)** Experimental design of pup separation test to examine changes in brain tissue dopamine levels. **k)** Pup separation significantly increases dopamine levels from baseline in dHF and NAc tissues (two-way ANOVA, main effect of time (F(3,52) = 4.287, p = 0.0089), main effect of brain region (F(1,52) = 225.7, p < 0.0001; interaction (F3,52) = 1.086, p = 0.3717), with significant increases in dHF dopamine levels (Dunnett’s multiple comparisons test, *p < 0.05, **p < 0.01). N = 5-9 animals/time point. **l)** Chronic pup separations increases baseline dHF dopamine concentrations (two-way ANOVA, interaction (F(3,53) = 4.822, p = 0.0048; group (1, 53) = 1.876), p = 0.1708, time (3, 53) = 1.736, p = 0.1706; Tukey’s multiple comparisons test, *p < 0.05, #p = 0.06). N = 5-12 animals/time point. Error bars represent mean ± SEM.

Since our cell-type deconvolution analyses indicated downregulation of neuronal markers, we first assessed proportional differences across major cell-types. A Monte Carlo permutation test confirmed significant shifts in several populations beyond random variation, with GABA.2 and GABA.4 clusters being reduced in Control PP dHF compared to both NP and Stress PP (**Fig. 4b-c**). We next used pseudobulk aggregation and differential expression analysis to identify DEGs within each annotated cluster (p < 0.05 and logFC > |1.5|). We observed transcriptional changes across both neuronal and non-neuronal subtypes, with excitatory neuron alterations predominantly observed between NP *vs.* Control PP, particularly within the subiculum (**Fig. 4d**). This pattern aligns nicely with enhanced contextual fear conditioning, as prior studies have implicated altered dHF excitatory outputs in improved learning outcomes in dams^24^. In contrast, the most pronounced neuronal changes between Control and Stress PP were observed in the GABA.2 and mossy cell populations, alongside widespread DEGs across non-neuronal cell-types (**Fig. 4d**); note that differential expression analysis could not be performed for the GABA.4 cluster due to insufficient sample representation in multiple groups. GO term analysis of DEGs from cell-types displaying proportional differences revealed significant enrichment in pathways associated with responses to endogenous stimuli (GABA.2, GABA.1, GABA.3, OPC), cell proliferation (GABA.2, CA1, immune cells), synaptic function (GABA.3, subiculum), and metabolic processes (Oligo.2, GABA.1) across both comparisons (**Fig. 4e-f**).

Probing deeper into these altered cell-types, we focused on the GABA.2 cluster owing to shared proportional and transcriptional changes observed in comparison to both NP and Stress PP groups, which suggested a common mechanism limiting dHF plasticity. Marker gene analysis revealed significant enrichment of dopamine receptor D1 *(Drd1)* and D2 *(Drd2)* expression within this cluster compared to other cell-types (**Extended Data 4j**), aligning with our prior bioinformatic analyses implicating dopaminergic regulation. To further resolve this population, we subclustered the GABA.2 cells and identified distinct subpopulations expressing *Drd1* and *Drd2*, both of which appeared to be altered in Control PP animals (**Fig. 4g**). In a separate cohort of mice, we validated these changes using RNA fluorescence in situ hybridization (FISH) in dorsal CA1 (**Fig. 4h-i, Extended Data 4k-l**), where dopamine receptor-expressing interneurons have been previously described^38,39^. While dopaminoceptive inhibitory neurons were implicated in our data, we examined whether such changes extend to other dopamine-sensitive neuronal populations. RNA FISH in the dentate gyrus, which contains excitatory *Drd1* and *Drd2*-expressing neurons (in granule and hilar mossy cells, respectively; **Extended Data 7a**), revealed significant group differences, alongside expected regional variations consistent with the sparser distribution of dopamine receptor-expressing neurons in CA1^40^ **(Fig. 4h-i, Extended Data 4m-n).**

Since these findings established dopaminergic signaling as a potential key mediator of persistent parity-induced adaptations, this prompted us to examine whether maternal stress disrupts this process within its defined postpartum window. Given prior evidence that acute pup separation elevates dopamine levels in NAc^41,42^, we first investigated whether a similar response occurs in dHF. Brain tissues were collected from 10 dpp dams at baseline, 30-minutes, and 3-hours post-separation, as well as 30-minutes post-reunion (**Fig. 4j).** We observed robust increases in dopamine levels in both NAc and dHF during separation, with expected regional differences reflecting the strength of innervation to these regions (**Fig. 4k**). Since acute and chronic stress produce different biological responses and behavioral outcomes, we next assessed the impact of repeated separation stress on dopamine modulation in these brain regions. Indeed, our data revealed distinct dopaminergic responses following acute *vs.* chronic separations, with repeated stress leading to loss of dopamine increases observed in response to pup separation, as well as elevated baseline dopamine in both dHF and NAc tissues (**Fig. 4l, Extended Data 4o**). Loss of induction following chronic stress may stem from elevated baseline levels, which facilitate engagement of autoreceptors and negative feedback mechanisms^43^. Similarly, increased release of dopamine in dHF tissues was not observed in NP females 30-minutes following separation from a littermate (**Extended Data S4p**), though this likely lacks the same salience as pup separation. These findings align with our snRNA-seq data, where decreased *Drd1/2*-expressing neurons in Control PP may reflect an adaptation to reduced dopamine-related transcriptional activity (Figs. 2 and 3). In contrast, elevated receptor expression in NP and Stress PP animals may represent a compensatory response to higher basal dopamine levels, as suggested by our observations following separation stress.

### Dopamine-dependent H3 dopaminylation enrichment is altered by parity in dHF of mice and humans

Together, changes in dopamine modulation and gene expression in our mouse model point to a role for dopamine-dependent epigenetic mechanisms in shaping persistent transcriptional states. Motivated by this, we turned to a recently characterized class of histone post-translational modifications dependent on intracellular pools of biogenic monoamines, including serotonin, dopamine, and histamine, termed monoaminylations^44–46^. This process involves transamidation of monoamines onto the glutamine 5 residue of histone H3 (H3Q5), a site adjacent to the canonically permissive lysine 4 tri-methylation (H3K4me3), by tissue transglutaminase 2 (TG2)^47^. Monoaminylation at H3Q5 can coexist with H3K4me3 and is responsive to physiological and environmental stimuli such as stress or drugs of abuse, with downstream effects on transcriptional output^44–46,48–52^. Using our previously validated H3K4me3Q5dop antibody^45^, we performed CUT&RUN-seq to test the involvement of H3 dopaminylation in parity transcriptional adaptations; note that this signal is lost in the absence of TG2 expression (**Extended Data Fig. 5a-c**). Following peak calling, we found that the majority of H3K4me3Q5dop peaks annotated to genic loci, with the majority of these peaks occurring within 2 kB of transcriptional start sites (TSS; **Extended Data 5d-g**). In all groups, we observed that gene expression varied significantly across quartiles of H3K4me3Q5dop enrichment, with higher signal associated with increased gene expression (**Extended Data Fig. 5h-i**).

To examine group-specific changes in H3 dopaminylation, we identified differentially enriched peaks using the DiffBind pipeline^53^. Notably, the majority of differential peaks were downregulated in Control PP dHF compared to both NP and Stress PP groups (**Extended Data Fig. 5j, Extended Data Tables 7-8**), consistent with prior results suggestive of reduced dopamine tone. Integrating these peaks with DEGs from the same comparisons revealed significant overlap between downregulated peaks and genes decreased in Control PP (**Extended Data Fig. 5k**), implicating loss of H3 dopaminylation in the downregulation of genes modulated by parity. To gain more resolution, we assessed H3K4me3Q5dop enrichment specifically at the TSS of parity-regulated DEGs. Again, Control PP dHF exhibited significantly reduced signal at most loci compared to NP and Stress PP (**Fig. 5a–b, Extended Data Fig. 5l**). Stratifying DEGs by directionality uncovered distinct transcription factors: downregulated genes were enriched for canonical repressors (Mbd3, Rest, Suz12), while upregulated genes were associated with ligand-responsive nuclear receptors (Esr1, Lxr, Ppara), pointing to ligand-dependent transcriptional programs that may compensate for reduced H3K4me3Q5dop by engaging alternate chromatin remodeling pathways (**Fig. 5c–d**).

**Figure 5.**
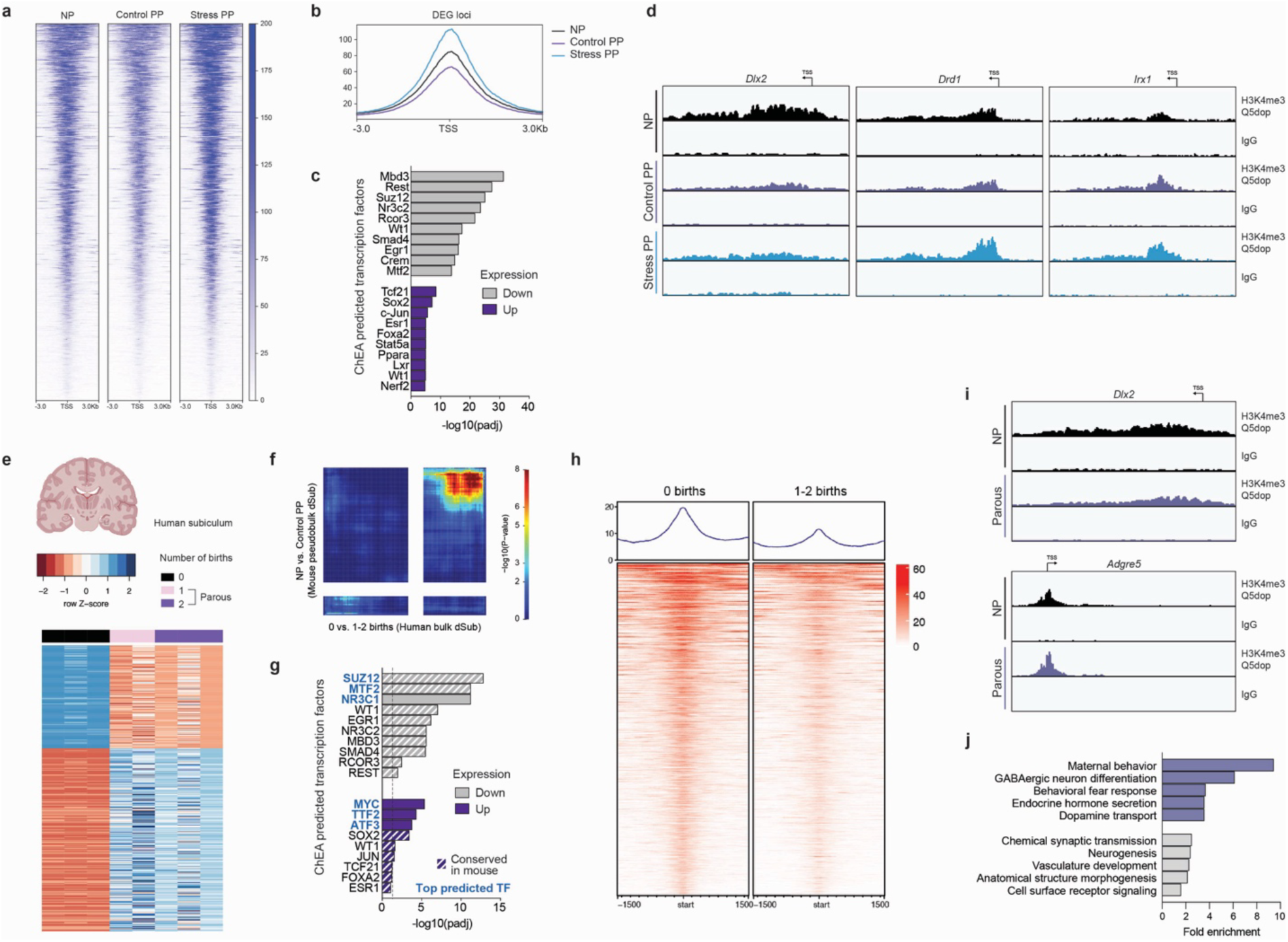
Parity downregulates H3 dopaminylation enrichment in dHF in both human and mice. **a)** Heatmaps and **(b)** profiles for H3K4me3Q5dop CUT&RUN-seq enrichment for NP vs. Control PP DEGs at TSS. Plots represent merged data from N=4 animals/group. **c)** ChEA ontology analysis identified significantly enriched transcription factors (TFs; adjusted *p* < 0.05) associated with downregulated DEGs, including a greater prevalence of repressive chromatin readers compared to those predicted for upregulated DEGs. **d)** Representative genome browser tracks for differentially enriched H3K4me3Q5dop (*vs.* IgG) peaks. Each track represents merged signal for N=4 samples/group. **e)** Heatmap of differential expression profiles of top 500 genes, by ascending p-values, from bulk RNA-seq of human dorsal subiculum (dSub) brain tissue, between NP subjects experiencing 0 births (N=3) *vs.* parous subjects previously having 1-2 births (N=5). **f)** Threshold-free comparison of human dSub RNA-seq (0 *vs.* 1-2 births) and mouse dSub pseudobulk from snRNA-seq (NP *vs.* Control PP) by rank-rank hypergeometric overlap. Pixels represent the overlap of differential expression profiles indicated, with color representing extent of significance. The lower left and upper right quadrants represent concordant gene regulation. **g)** ChEA ontology analysis showing the top three significantly enriched TFs (blue font; adjusted p < 0.05) associated with downregulated and upregulated DEGs from human dSub RNA-seq data. Additional TFs conserved between human and mouse analyses are indicated by white stripes. The dotted line indicates the threshold of 1.3. **h)** Heatmaps and profiles for H3K4me3Q5dop CUT&RUN-seq enrichment for human dSub, comparing subjects with 0 *vs.* 1-2 births at differential loci (*p* < 0.05, log_2_FoldChange ≥ |0.1|). Plots represent merged data from 3 NP and 5 parous subjects. **i)** Representative genome browser tracks for differentially enriched H3K4me3Q5dop (*vs.* IgG) peaks from human dSub. Each track represents merged signal for N=3-5 samples/group. **j)** GO ontology analysis of differential H3K4me3Q5dop loci identified top significantly enriched pathways (gray bars; ranked by adjusted *p*) related to synaptic signaling, neurogenesis, and vasculature development. Additional significant pathways linked to maternal behavior, fear response, and dopamine transport are shown in purple.

To assess the translational relevance of these findings, we next performed transcriptomic and H3K4me3Q5dop profiling in human dorsal subiculum (dSub), a subregion of the dHF that serves as its primary output and was identified in our snRNA-seq analysis as the most transcriptionally responsive subregion to parity. Comparing age-matched NP subjects with parous subjects who had 1-2 previous births (**Extended Data Table 9**), we identified robust differential gene expression indicative of long-lasting effects of parity in human brain (**Fig. 5e**). These transcriptional signatures showed strong concordance with those observed in our mouse model, as determined by comparison to NP *vs.* Control PP dSub pseudobulk expression profiles from our snRNA-seq data, particularly for downregulated genes (**Fig. 5f**), and similarly displayed enrichment in pathways related to extracellular matrix remodeling and metabolism (**Extended Data Fig 6a**). Furthermore, predicted upstream transcription factor ontologies were conserved across species, with both upregulated and downregulated DEGs in human dSub enriched for regulators similarly identified in mouse (**Fig. 5g**). We also performed H3K4me3Q5dop CUT&RUN-seq in human dSub tissues to associate such persistent gene expression signatures with dopamine-dependent chromatin remodeling. Similar to our mouse tissues, we observed that the majority of H3K4me3Q5dop peaks occurred at genic loci (**Extended Data Fig. 6b-d**), with increasing signal corresponding to increased gene expression (**Extended Data Fig. 6e-f**). Next, we performed differential binding analysis comparing parous and NP subjects. Consistent with our mouse findings, parous dSub exhibited a widespread reduction in dopaminylation signal that significantly corresponded with our DEG analysis (**Fig. 5h-i; Extended Data Fig. 6g; Extended Data Table 10**). Pathway analysis of differentially enriched peaks revealed significant enrichment for loci involved in maternal behavior, behavioral fear responses, hormone regulation, and dopamine transport - processes similarly implicated in our mouse model (**Fig. 5j**). These findings suggest that parity induces long-lasting dopamine-dependent remodeling of epigenomic programs in a region of the human dHF, mirroring mechanisms observed in rodents.

### Chemogenetic suppression of dopamine release into dHF recapitulates key features of persistent parity-induced plasticity

Together, these findings point to conserved, parity-associated remodeling of dopamine-sensitive transcriptional programs in dHF. We therefore hypothesized that persistent reductions in dopamine tone may engage adaptive chromatin mechanisms to support enduring transcriptional and behavioral plasticity. To assess this, we chemogenetically suppressed the dopaminergic VTA projection to dHF by bilaterally expressing a floxed inhibitory neuronal hM4Di-DREADD fused to mCherry (*vs.* floxed mCherry controls) in the VTA, coupled with retrograde AAVs expressing a tyrosine hydroxylase (TH)-dependent Cre recombinase into the dHF (**Fig. 6a**). Selection of the VTA was guided by our findings that pup separation stress drives coordinated regulation in the dHF and NAc, along with its established role as a key upstream modulator of dopaminergic signaling in the maternal NAc^54^ (also supported by our data; **Extended Data 4q**). We validated this approach by confirming selective mCherry expression in TH+ neurons of the VTA, as well as reduced *Fos* expression following administration of the selective DREADD agonist deschloroclozapine (DCZ; **Fig. 6b, Extended Data 5a-b**.) To minimize injection-related stress in this paradigm, we sought to restrict the window of chronic DCZ administrations. Since maternal stress from 10–20 dpp was sufficient to disrupt parity-induced plasticity, we posited that dHF dopamine suppression during this period may correspondingly be sufficient to promote neuronal adaptations in virgin NP (NP-mCherry, NP-hM4Di; **Fig. 6c**). As our comparisons aimed to determine if chronic dopamine reduction in virgin females can phenocopy parity-induced adaptations, we administered DCZ to postpartum dams in parallel (PP-mCherry, PP-hM4Di).

**Figure 6.**
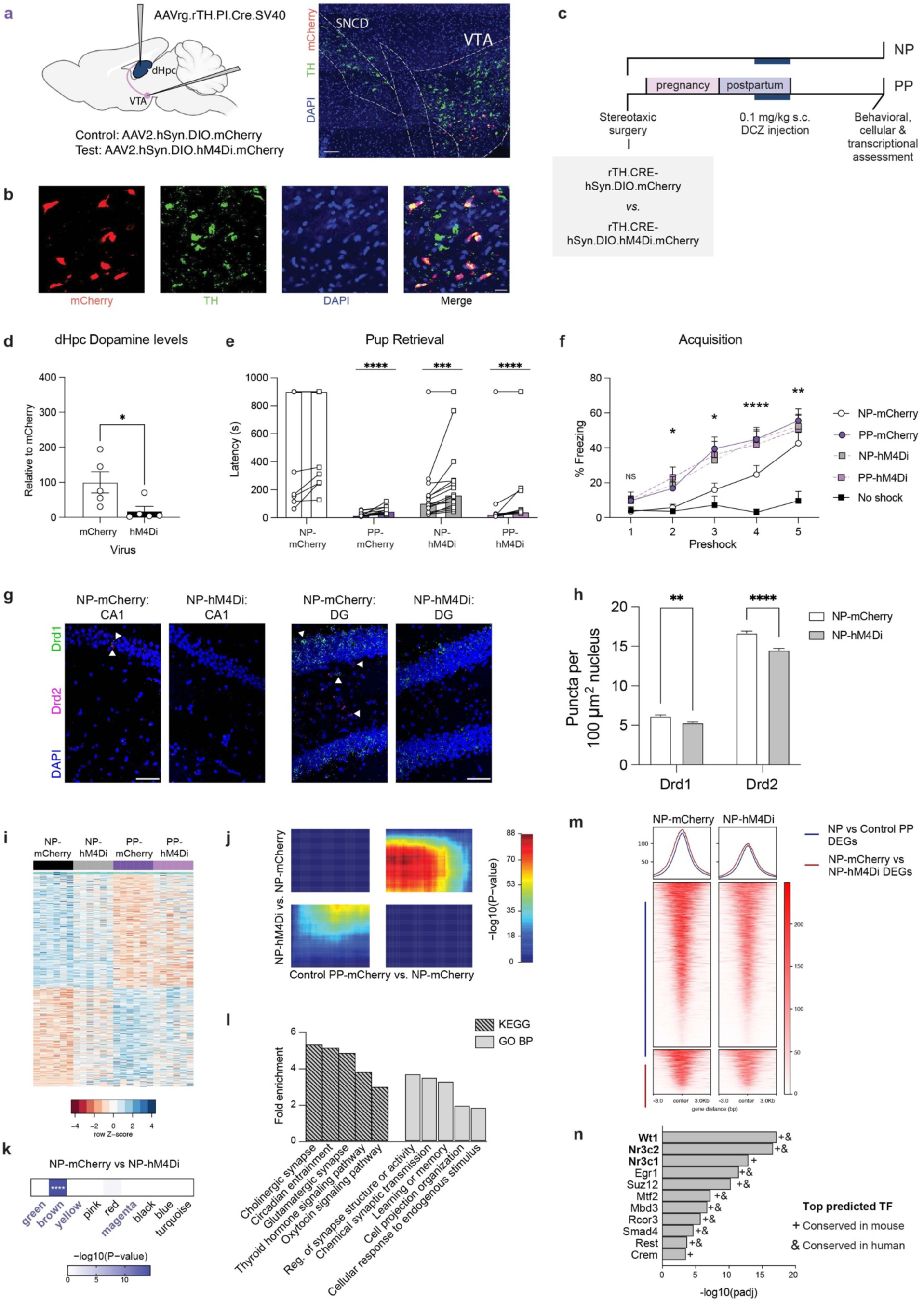
Chronic dopamine suppression is sufficient to mimic persistent maternal dHF plasticity. **a)** Experimental design (left) and representative image (right) of AAV2.hSyn.DIO.mCherry targeting to VTA projection neurons following dHF injection of retrograde (rg) Cre under control of the tyrosine hydroxylase (TH)’. Scale bar, 100 μm. **b)** Representative images showing overlap of virally-induced mCherry with TH+ cells in VTA. Scale bar, 20 μm. **c)** Timeline of stereotaxic surgeries and s.c. deschloroclozapine (DCZ) injections from 10-20 dpp in NP and PP female mice, followed by experimental assessments starting at 49 dpp. **d)** Significant reduction in dHF dopamine levels following inhibition of VTA projection neurons (Student’s t-test:t(8) = 2.477, *p = 0.0383). N=5/group. **e)** PP-mCherry, NP-hM4Di, and PP-hM4Di retrieved pups faster than NP-mCherry (two-way rmANOVA, main effect of group (F(3,47) = 14.39, p < 0.0001), main effect of pup number (F(1, 47) = 18.96, p < 0.0001; Sidak’s multiple comparisons test, ****p < 0.0001, ***p < 0.001). N =11-15/group. Bar represents mean. **f)** PP-mCherry, NP-hM4Di, and PP-hM4Di freeze more compared to no shock controls at earlier phases of contextual fear acquisition, compared to NP-mCherry (two-way rmANOVA, main effect of group (F(4,41) = 4.739, p = 0.0031), main effect of preshock (F(2.818,115.5) = 61.92, p < 0.0001), interaction (F(16,164) = 2.606, p = 0.0012; Tukey’s multiple comparisons test, *p ≤ 0.05, **p < 0.01, ****p < 0.0001). N = 9-12/group. Error bars represent mean ± SEM. **g)** Representative images of RNAScope for *Drd1* and *Drd2* mRNAs in dorsal CA1 (top) and DG (bottom). Scale bars, 50 μm. **h)** Quantification of *Drd1* and *Drd2* mRNA puncta in CA1 and DG nuclei (two-way ANOVA, main effect of group (F(2,33404) = 64.43, p < 0.0001), main effect of gene (F(1,33404) = 2770, p < 0.0001), interaction (F(2,33404) = 12.52, p = 0.0004); Sidak’s multiple comparisons test, **p < 0.01, ****p < 0.0001). N = 3 animals/group. **i)** Bulk dHF differential expression profiles of top 500 genes (by ascending p-values) between NP-mCherry and PP-mCherry. N=6/group. **j)** Threshold-free comparison by rank-rank hypergeometric overlap, showing concordant gene regulation (bottom left and upper right quadrants) between the comparisons indicated. **k)** Overlap of DEGs (NP-hM4Di vs. NP-mCherry, adj. p < 0.05) with gene co-expression modules (****p < 0.0001). High parity sensitivity modules are indicated in bold. **l)** Select GO terms and KEGG pathways significantly enriched for NP-mCherry vs. NP-hM4Di DEGs (FDR < 0.05). **m)** Heatmaps and profiles for H3K4me3Q5dop CUT&RUN-seq enrichment for NP-mCherry vs. NP-hM4Di at TSS for DEGs between NP vs. Control PP (top, blue line) and NP-mCherry vs. NP-hM4Di (bottom, red line). Plots represent merged data from N=4 animals/group. **n)** ChEA ontology analysis of downregulated DEGs identified significantly associated TFs (adjusted *p* < 0.05), paralleling patterns observed with parity in both human and mouse datasets.

After confirming that our approach effectively suppressed dHF dopamine (**Fig. 6d**) – and did not induce gross metabolic changes (**Extended Data 5c**) – we assessed behavioral outcomes 1-month after the final DCZ injection (49 dpp). In the pup retrieval test, NP-hM4Di females exhibited significantly reduced latency to retrieve pups compared to NP-mCherry, with no differences observed relative to PP dams (**Fig. 6e**). Notably, while 8/13 NP-mCherry females failed to retrieve either pup, this occurred in only 1/15 NP-hM4Di animals, comparable to PP-mCherry (0/12) and PP-hM4di (1/11) dams. In contextual fear conditioning, NP-hM4Di females demonstrated enhanced acquisition, mirroring the heightened learning observed in PP dams (**Fig. 6f**). In the context recall test, NP-hM4Di females showed no significant differences from PP dams (**Extended Data 5d**), though subtle effects were observed, suggesting that additional neuromodulatory mechanisms may interact with dopamine to modulate the consolidation and retrieval of conditioned fear responses underlying maternal behavioral plasticity.

Next, we examined whether such dopamine suppression influences cellular receptor expression in the dHF. Similar to Control PP, NP-hM4Di females exhibited sustained downregulation of *Drd1* and *Drd2* in both CA1 and dentate gyrus (**Fig. 6g-h, Extended Data 5e-h**). Next, we conducted transcriptional profiling of the dHF in virally infected groups to assess the degree to which dopamine suppression mirrors parity-associated gene expression patterns. Comparing NP-mCherry *vs.* PP-mCherry expression profiles from bulk RNA-seq data, we observed an intermediate expression pattern in NP-hM4Di dHF tissues (**Fig. 6i, Extended Data Table 11**). RRHO analysis further supported transcriptional concordance between PP-mCherry and NP-hM4Di groups in comparison to NP-mCherry (**Fig. 6j**). To next assess whether the long-term transcriptional effects of prior baseline dopamine suppression align with changes in parity-associated transcriptional programs, we compared NP-mCherry *vs.* NP-hM4Di DEGs to previously identified parity-sensitive modules. Consistent with our behavioral findings, DEGs from NP-hM4Di overlapped with the brown module but not other parity-sensitive modules (**Fig. 6k**), thereby highlighting a specific mechanistic contributor to behavioral adaptations within the broader transcriptional and neuromodulatory landscape shaped by parity. Functional annotation of these DEGs revealed enrichment in pathways governing synaptic plasticity and responses to endogenous stimuli (**Fig. 6l**), which mirrored those found to be altered in Control PP dHF.

Finally, to assess whether the effects of dopamine suppression on dHF transcription are modulated in part through H3 dopaminylation, we conducted CUT&RUN-seq targeting H3K4me3Q5dop in virally-infected NP tissues. Examining H3K4me3Q5dop enrichment at the TSS of both parity-regulated DEGs and genes altered between NP-mCherry and NP-hM4Di groups, we observed significantly reduced signal at all loci examined in NP-hM4Di tissues (**Fig. 6m; Extended Data Fig. 7i-j**). We identified significant transcription factors predicted for downregulated DEGs between NP-mCherry and NP-hM4Di (none for upregulated DEGs), overlapping with those observed in parous subjects in both human dSub and our mouse model (**Fig. 6n**). Overall, these data demonstrate dopaminergic modulation as a key mechanism underlying the molecular and functional adaptations associated with parity in the maternal brain (**Extended Data 5k**).

## DISCUSSION

Despite decades of evidence that matrescence induces long-lasting behavioral adaptations across species, the mechanisms underlying these phenomena remain poorly understood. Through controlled brain-wide, time-course, and cell type-specific transcriptomic analyses – paired with robust behavioral outputs, we identified gene networks, reproductive events, and neuromodulatory pathways that dynamically shape regional sensitivity across pregnancy and postpartum experiences, thereby promoting plasticity long after these stages have ended. In particular, our study identified the dHF as a key site of sustained plasticity, consistent with its established roles in spatial cognition, novelty detection, and sensory integration, which may bestow parous dams with heightened sensitivity to salient environmental cues relevant for resource foraging, nest navigation, and offspring survival^55–58^. While not traditionally considered a maternal behavior-regulating region, the dHF integrates these salient cues to shape pup retrieval responses. Consistent with our findings, lesion studies corroborate hippocampal involvement in pup retrieval behavior^59,60^. Moreover, hippocampal activity modulates downstream circuits via the polysynaptic VTA-hippocampal loop, which projects to the NAc and ventral pallidum - key regulators of pup retrieval^58,61^. Notably, recent human studies show that maternal early life stress alters hippocampal responses to infant cues^62,63^, supporting chronic stress as a common interferer in the normal trajectory of maternal dHF neuroadaptations. Building on this, our maternal-pup separation stress model revealed that dopamine dynamics are essential for maintaining long-term neural and behavioral adaptations in the maternal dHF. Indeed, in the current study we established the necessity and sufficiency of maintaining appropriate dopamine tone for parity-related dHF plasticity, demonstrating that chronic dopamine elevation through maternal stress, and chemogenetic suppression within virgin dHF, bidirectionally modulate parity-related changes at transcriptional, epigenetic, behavioral, and cellular levels. Moreover, we establish conserved transcriptional and epigenetic changes in the human dSub, a subregion of the dHF. Such findings implicate dopamine as a key driver of long-term maternal brain remodeling, expanding upon the prevailing view that maternal dopamine function is solely driven by transient surges during maternal behaviors, instead revealing a concurrent downregulation of dopamine tone in critical regions, such as the dHF, to modulate persistent behavioral and cellular plasticity.

While our study focused largely on dopamine, it is notable that our chemogenetic manipulations recapitulated key transcriptional and behavioral outcomes of parity, albeit not to the same extent. This underscores dopamine as a crucial but not exclusive driver of parity-induced adaptations. Indeed, our data also implicate hormonal contributions, including estrogen, progesterone, and oxytocin signaling, in shaping transcriptional programming. Consistent with this, previous studies have linked parity to estradiol-driven neuronal alterations^12^, while others have demonstrated that blocking oxytocin signaling disrupts parity-induced spatial learning enhancements^27^. These findings suggest that parity programming arises from integration of multiple neuromodulatory pathways to induce long-term effects. However, given that estrogen signaling during postpartum regulates oxytocin release, which in turn modulates VTA signaling, we propose that dopamine adaptations serve as a key downstream mechanism within a broader network driving sustained maternal neuroplasticity^64^. Together, this work demonstrates that transient neuromodulatory processes during pregnancy and postpartum drive lasting maternal neuroplasticity. Importantly, this fundamental shift in brain state may interact with future experience-dependent plasticity. As parity is implicated in both risk and resilience to brain disorders^65^, future research should account for parity status as a key variable shaping differential outcomes, particularly in its interactions with other risk factors. In particular, our findings highlight an interplay between parity and stress, a key risk factor for brain disorder vulnerability, in shaping maternal brain outcomes. Thus, this study provides novel insights into the gene networks, dynamics, and neuromodulatory pathways that drive long-lasting parity-induced adaptations, while underscoring the importance of stress mitigation during pregnancy and postpartum.

## ACKNOWLEDGEMENTS

We would like to thank members of the Maze lab for their helpful discussion on this study, and members of the Nestler, Russo, and Kenny labs at the Icahn School of Medicine at Mount Sinai for providing materials and/or access to behavioral equipment. We thank Dr. Min Chen, Winnie Chen, Emma Andraka, and Rasika Iyer for experimental assistance. This work was partially supported by grants from the National Institutes of Health: R01 MH116900 (I.M.), F32 MH126534 (J.C.C.), F31 NS132558 (A.M.C.), F99 NS139541 (A.M.C.), as well as funds from the Brain and Behavior Research Foundation (J.C.C.), and the Howard Hughes Medical Institute (I.M.). All schematics were created with Biorender.com.

## AUTHOR CONTRIBUTIONS

J.C.C. and I.M. conceived of the study, designed the experiments, and interpreted the data. J.C.C., G.D.S., A.M.C., and S.D. performed behavioral experiments. J.C.C., G.D.S., A.M.C., and S.D. performed surgeries. J.C.C., G.D.S., and C.Z. performed estrous cycle analyses. J.C.C., G.D.S., and S.D. conducted immunofluorescence and RNAscope analyses. J.C.C., G.D.S., and B.H.W. performed molecular experiments. J.C.C. and E.A.B. performed single-cell sequencing experiments. J.C.C., G.D.S., E.W. and B.H.W. performed bioinformatic analyses. N.M. and G.T. provided human tissues. J.C.C. and I.M. wrote the manuscript.

## SUPPLEMENTARY MATERIALS

Materials and Methods

Extended Data Figs. 1 to 7

Extended Data Tables 1 to 9

## MATERIALS AND METHODS

### Human subjects

Brain tissue used in this study was provided by the Douglas-Bell Canada Brain Bank (DBCBB; www.douglasbrainbank.ca; RRID:SCR_025991). Informed consent from next-of-kin was obtained for each individual included in this study. Psychological autopsies, considered the gold standard for obtaining information on deceased individuals^66,67^, were conducted. Briefly, these consist of a series of proxy-based, structured interviews assessing psychopathology with next-of-kin and complemented by reviews of medical records, as previously described^66^. Groups were matched for depression diagnosis, and were otherwise neurotypical individuals who died suddenly without prolonged agonal periods and did not have evidence of axis I disorders. Groups were matched for postmortem interval (PMI), tissue pH, and RNA Integrity number. Frozen histological grade samples of gray and white matter were dissected from the subiculum by expert neuroanatomists and stored at –80 °C. Dissections were performed on 0.5 cm-thick coronal sections with the guidance of a human brain atlas^68^ (see also http://www.thehumanbrain.info/brain/bn_brain_atlas/brain.html). Subiculum samples were obtained from sections equivalent to plate 43 of the atlas (level of lateral geniculate nucleus), by dissecting through the hippocampal fissure with a slight upward angle, up to the beginning of the CA1 region.

### Animals

Wild-type C57BL6/J mice were purchased from Jackson Laboratories at 8-weeks old, and maintained on a 12-h/12-h light/dark cycle throughout the entirety of the experiments. Mice were provided with *ad libitum* access to water and food throughout the entirety of the experiments. All behavioral testing occurred during the animals’ light cycle. Experimenters were blind to experimental group, and the order of testing was counterbalanced during behavioral experiments. All animal procedures were performed in accordance with NIH guidelines and with the approval of the Institutional Animal Care and Use Committee of the Icahn School of Medicine at Mount Sinai.

### Breeding

Adult virgin female mice were pair bred in-house with age-matched males. Males were removed after a maximum of 5 days, and pregnant females were singly-housed at least 2 days before parturition. Pups were counted on the day of birth (0 dpp) and weaned at 21 dpp. Only dams with litters between 4-10 pups were used for all experiments. On the day of weaning, dams were group-housed into cages of 3-5 mice with animals of the same experimental condition. For timed breedings, copulation plugs were checked every morning within 1-hour after lights on, where confirmation of a plug was designated as E0.5, signaling the immediate removal of the female to her own cage with a nestlet. Virgin NP females were age-matched for each experimental cohort. Mating-experienced NP females were confirmed for the presence of a copulation plug. For Pregnancy Only dams, pups were removed at 0 dpp within 3 hours after lights-on to minimize maternal-offspring interactions. For comparison of PP dams to pup sensitized virgin females, litters were culled to 4 pups to equate litter size with the number of pups used for each sensitized female. In all other experiments, litters were not culled.

### Postpartum stress paradigm

Stress PP females were subjected to limited nesting and maternal separation from 10-20 dpp, as previously described^69–71^. On each day of separation, the entire litter was removed to a clean cage with Sani-Chip bedding for 3-4 hours. Separations occurred during the light cycle, and the timing varied each day to minimize predictability and acclimatization. EnviroDri nesting material was depleted to 1/3 of control cages during the days of separation. Following pup weaning on 21 dpp, the nesting material was restored to normal levels, and dams were group-housed into cages of 3-5 with animals of the same experimental condition.

### Pup sensitization

Pup sensitization was conducted as previously described^22,72^. On the first day of pup sensitization, virgin NP females were presented with four pups (postnatal day 5) from a cage consisting of a lactating donor dam and her litter. Donor pups were exchanged for satiated pups from the same litter every 8-12 hours for 21 days. Donor pups were weighed daily to ensure continual weight-gain throughout the experiment, and only pups that exhibited consistent weight-gain were used. Behavioral observations were conducted during the first four days of sensitization. Each dam was observed for 30 minutes during the light phase for the following maternal behaviors: licking/grooming, crouching, and nestbuilding. During these observation periods, females were closely monitored to ensure that they did not display aggressive behavior toward the donor pups. On the last day of sensitization, donor pups were weaned from the cage, and sensitized females were group-housed into cages of 3-5 mice with animals of the same experimental condition.

### Brain tissue collections

Animals were sacrificed by rapid decapitation. Whole brains were flash frozen with cold 2-methylbutane and stored at −80°C until further processing. Flash-frozen brains were sectioned at −20°C using a 1 mm mouse coronal brain matrix (Stoelting Co.). Tissues enriched for the brain region of interest were micropunched using a hollow needle (Ted Pella) according to the Allen Brain Atlas (see **Fig. S1A**).

### Behavioral analyses

All animals (18-30 weeks, depending on the time from pup weaning) were handled for 2 minutes for two consecutive days prior to initial behavioral testing. Animals were habituated to the testing room for 1 hour prior to each behavioral assay. All testing occurred during the light phase.

#### Pup retrieval

All animals were individually housed for 24-hours prior to testing. Following habituation, 2 pups from a donor litter with a lactating dam (aged 4-6 dpp) were placed in different corners opposite to the nest in the home cage. Pup-directed interactions were recorded from above for 15-minutes or until both pups were successfully retrieved into the nest. Retrieval latency was calculated as [time pup was first placed into the nest by animal - time first pup was placed into the cage by experimenter].

#### Contextual fear conditioning

On day 1, mice were habituated for 10 minutes to the testing chamber, which consisted of a square plexiglass box with a metal grid inside a sound-attenuating cabinet wiped with 70% ethanol (Med Associates, VT). On day 2, following a 3-minute baseline measurement, mice were trained with five 2.0-second 0.7 mA foot shocks delivered with an intertrial interval of 90-seconds. Testing for conditioned fear responses (freezing) for a total of 5-minutes occurred 24-hours following training. Freezing was measured using ANY-maze software connected to a camera positioned above the testing chamber. Freezing is expressed as a percentage of the total test time or as a percentage of the 60-seconds prior to each shock during conditioning. Animals with freezing levels exceeding 40% prior to the first shock were excluded to remove potential confounding effects of heightened baseline responsivity that could interfere with accurately assessing conditioned fear.

#### Open Field

Mice were placed in a 16×16 square arena under dim lighting for 5 minutes. A camera positioned overhead recorded the total distance and time spent in the center *vs.* periphery using Ethovision software.

#### Object location task

For training, animals were allowed to freely explore two identical objects placed equidistant from adjacent corners of a 16×16 cm square arena for 5 or 10 minutes, before being returned to the home cage. Following a 1-hour delay, the animals were returned to the arena for testing, during which time one object was moved to the opposing corner. During the test, the animals were allowed to freely explore the objects for 5-minutes. A camera positioned overhead recorded time spent exploring each object using Ethovision software. The discrimination score was calculated as [(time spent with moved object) – (time spent with unmoved object)/[(time spent with moved object) + (time spent with unmoved object)].

#### Light-dark box

Anxiety-like behavior was tested in an apparatus containing two interconnected 20×20 cm compartments (Omnitech Electronics Inc.). One compartment was illuminated during the session (“light” side), while the other was covered by an opaque black perspex lid (“dark” side). The distance, time spent, and number of crossovers in each compartment were automatically recorded by Fusion software during the 10-minute test.

#### Forced Swim Test

Mice were placed in a 4 liter glass beaker with 2 liters of room temperature water for 8 minutes. Each session was recorded and scored by a blinded observer. The total number of seconds that mice were immobile during the last 5 minutes of the test were recorded, as previously described^73^.

#### Observation of pup-directed behaviors in the home cage

Confirmation of maternal behavior in pup sensitized females was conducted based on prior studies^74^. Animals were observed from the first to fourth day of pup sensitization, based on published work that maternal sensitization in C57BL6/J females occurs following 4-days of pup exposures^29,75^. Observations occurred in the light period in the first 30-minutes after pups were placed in the cage. Each animal was scored every 3-minutes, with the observed behavior recorded as one or more of the following categories: nestbuilding, grooming, sniffing, crouching, nursing, eating, drinking, self-grooming, no-contact. PP dams were scored for the first 4-days following birth (0-3 dpp) concurrently for comparison.

### Estrous cycle testing

Vaginal samples were taken on each day of behavioral testing. 15 µL of sterile PBS was gently pipetted into the vagina and mounted on a glass slide. Vaginal smears were stained with crystal violet dye, washed twice with water, and cover slipped with glycerol. Three 20x images per sample were acquired on a light microscope. Estrous stage was determined using the pretrained network of EstrousNet^76^, and confirmed afterwards by a trained experimenter based on cytology of nucleated, cornified, or leukocytic cells.

### Viral Transduction

Mice were anesthetized with ketamine (100mg/kg) and xylazine (10 mg/kg) i.p. and positioned in a stereotaxic frame (Kopf instruments). Given the widespread changes in dopamine receptor expression across dHpF subregions, we did not restrict our viral manipulations to a specific subregion. 2 µl of retrograde AAV.rTH.PI.Cre.SV40 (titer ≥ 7×10¹² vg/mL, Addgene #107788-AAVrg) was infused bilaterally into the dHpF at 0.2 uL/min using the following coordinates: 7° angle; anterior-posterior (AP) -2.2 mm, medial-lateral (ML) ±2.0 mm, dorsal-ventral (DV) -2.0 mm. 1 µl of pAAV-hSyn-DIO-mCherry (titer ≥ 4×10¹² vg/mL, Addgene #50459-AAV2) or pAAV-hSyn-DIO-hM4D(Gi)-mCherry (titer ≥ 5×10¹² vg/mL; Addgene #44362-AAV2) was bilaterally infused into the VTA using the following coordinates: 7° angle; AP -3.3 mm, ML ±0.9 mm; DV -4.6 mm. Needles remained in place for 7 minutes following injection to minimize virus diffusion. Viral validations were conducted at least 21 days post-surgery to allow for optimal viral expression and recovery.

### Chemogenetic manipulation

To directly manipulate dopamine signaling in dHpF, we selectively targeted the VTA-dHpF projection during a window established as being critical for dopaminergic downregulation. Following 7 days of recovery, surgerized female mice were randomly assigned to NP or PP groups. PP females were pair bred with naïve male mice for a maximum of 5 days, and individually housed prior to parturition. From 10-20 dpp, PP females were injected subcutaneously with deschloroclozapine (DCZ, Tocris #7193) – to minimize off-target effects^77^ - at 1ug/kg in 1% DMSO or vehicle (saline). NP females were injected with DCZ concurrently. Following weaning on 21 dpp, PP dams were group-housed with animals of the same experimental condition. Behavioral testing occurred beginning at 28-days post-weaning (49 dpp), and brain tissues were collected following contextual fear conditioning for further analyses.

### RNAscope in situ hybridization and analysis

Fresh frozen brains were cut into 12-16 μm thick slices in the coronal plane with a cryostat (Leica CM3050-S), mounted on charged Superfrost Plus microscope slides (Thermofisher #P36934), and stored at -80°C until processing. Sections were post-fixed with 4% PFA for 1.5 hours at 4°C, and permeabolized with hydrogen peroxide (10 minutes RT) and Protease III (30 minutes RT). The RNAscope Multiplex Fluorescent Reagent Kit v2 (ACD Bio) was used according to manufacturer’s instructions to sequentially stain sections with the following probes: Drd1a (Mm-Drd1a-C1, #461901) and Drd2 (Mm-Drd2-C3, #406501-C3). RNAscope probes were visualized using TSA Vivid Fluorophores (Tocris) at 1:750 dilution. Sections were counterstained with DAPI (ACD Bio) and mounted using ProLong Gold Antifade Mountant (Thermo). Confocal images (3 images per animal, 1024 × 1024 pixels) were acquired on a Zeiss LSM 780 upright microscope using a 40X objective with Zen Black software, with 5×1 tiled images. Images were averaged across 8 consecutive acquisitions at a bit depth of 16 bits, with 2 z-stacks acquired per image. Subcellular quantification of individual puncta per 100 μm nucleus - identified by cellular detection of DAPI staining - was performed for each maximum intensity projected image using QuPath software (v.0.5.1)^78^. Regions of interest (CA1 and dentate gyrus) were annotated and determined by superimposing images onto the Allen Brain Atlas.

### Immunofluorescence and analysis

Mice were anesthetized with isoflurane and perfused with cold 1×PBS followed by 4% PFA. Brains were post-fixed in 4% PFA overnight and then transferred to a 30% sucrose/PBS solution for 2 days. Brains were sectioned at 40 µm thickness using a Leica CM3050-S cryostat, with serial sections collected from the VTA. For each subject, 2–3 brain slices were blocked for 2 hours (0.1% Triton X-100, 10% normal donkey serum), followed by overnight incubation at 4 °C with primary antibodies: chicken anti-tyrosine hydroxylase (1:500, Aves Labs #TYH) and rabbit anti-Fos (1:2000, Synaptic Systems #226-008). The next day, slices were incubated for 2 hours at RT with fluorescent-conjugated secondary antibodies (donkey anti-chicken Alexa Fluor 488, Invitrogen #A78948; donkey anti-goat Alexa Fluor 680, Invitrogen #A10043). Slices were counterstained with DAPI (1:10000, Thermo Scientific #62248) and mounted with ProLong Gold Antifade Mountant (Thermo Fisher #P36934). Confocal images (2-3 replicates per animal, 1024 x 1024 pixels) were acquired on a Zeiss LSM 780 upright microscope using a 40x objective with Zen Black software. Images were averaged across 8 consecutive acquisitions at a bit depth of 16 bits. Image analysis was conducted using FIJI software (NIH). The Fos and TH channels were thresholded using the MaxEntropy method to define regions of interest. Following identification of colocalized Fos+/TH+ signal, Fos intensity was measured and averaged from 4-6 images per animal. Brightness and contast were adjusted for representative images.

### Dopamine ELISA

Brain tissue dopamine levels were assessed in response to 3 hours of pup separation in the home cage. Independent biological replicates were collected at each time point, with separate animals used for each measurement. Brains were rapidly harvested (see Brain Tissue Collection, above) at baseline (prior to pup removal, 0 min), during pup removal (30 and 180 min), and 30 min after pups were returned to the home cage (210 min). For assessment of NP females, samples were collected in the home cage 30-min following removal of a littermate. Equal weights of micropunched brain tissues were homogenized in lysis buffer (0.01 N HCl, 1 mM EDTA, 4 mM sodium metabisulfite). Tissue dopamine levels were assessed using the Dopamine (Research) ELISA Kit (ALPCO Diagnostics) according to manufacturer’s instruction.

### HPA axis assessment

Plasma corticosterone levels were assessed in response to 3-hours of pup separation in the home cage. Testing was initiated within 2-hours after lights on. Tail blood was collected prior to pup removal (0 min), during pup removal (30 and 180 min), and 120 min after pups were returned to the cage (300 min) from the same animals. Samples for control animals in the home cage, used to account for the effects of handling-induced stress, were collected concurrently. Blood samples were immediately mixed with 50 mM EDTA and centrifuged at 5000 rpm for 10 minutes. Plasma was collected and stored at −80 °C until analysis. Corticosterone levels were quantified using a Corticosterone ELISA kit (ENZO Life Sciences) according to manufacturer’s instruction.

### Clonal *TGM2* Knockout in HeLa cells

HeLa cells (ATCC, CCL-2) were cultured at 37 °C with 5% CO_2_ in DMEM medium (high-glucose, ThermoFisher 11965118) supplemented with 10% FBS (Sigma-Aldrich) and 500 U ml−1 penicillin and streptomycin. The CRISPR guide RNA targeting exon 5 of *TGM2* was purchase from IDT, containing the sequence ACGCTGGGACAACAACTACG. 120 pmol of guide RNA and 100 pmol of Alt-R™ S.p. Cas9 Nuclease V3 (IDT, 1081058) were premixed for 15 minutes in 5 μL total (with PBS). 200k HeLa cells were washed in PBS, before resuspending in 20 μL of nucleofector solution (SE Cell Line 4D-Nucleofector® X Kit S, Lonza V4XC-1032). The 5 μL Cas9/sgRNA mix was added, as well as 1 μL of Alt-R Cas9 Electroporation Enhancer (IDT, NC1395977), and all mixed gently. The entire mixture was transferred to a 16-well Nucleocuvette strip (Lonza, PDH-2104) gently, and nucleofected using the Lonza 4D-Nucleofector® X Unit, using the default settings for HeLa cells. Directly after nucleofection, 80 uL of pre-warmed culture media was added to the cuvette. The entire mixture was immediately transferred to a 6 well plate with 2 mL of prewarmed culture media, and incubated cultured at 37 °C with 5% CO_2_ for 48 hours. Nucleofection efficiency was assessed by using a second positive control reaction with a pMax-GFP plasmid (Addgene, 177825). The pool of cells was diluted to a concentration of 1 cell per 200 uL, and 100 uL aliquoted into each well of 10 96 well plates, and cultured at 37 °C with 5% CO_2_. After 14 days, single-clones were identified using a light microscope. Single-clone containing wells were expanded and targeting assessed by extracting genomic DNA and performing PCR and PCR sequencing over the targeted site (Fwr: GGCTCCAGCCCCCACCATCTGCCGCAC, Rev: GCCACATAGCGCATTGAGAGTGTTGGT). PCR sequencing results were assessed using Synthego’s ICE analysis. Clones which were identified as introducing premature stop codons were expanded further.

### Assessment of *TGM2* KO Cell Line

#### Western Blot for TG2

Briefly, HeLa cells were collected and lysed using high-salt buffer (20 mM HEPES pH 7.9, 500 mM KCl, 10 mM MgCl2, and 1 % NP-40), followed by brief pulse-sonication. 100 ug of protein was run on a 4-12% Bis-Tris gel (Invitrogen, NW04122) for 45 minutes at 150V. Protein was transferred to a 0.2 µm nitrocellulose membrane using a Trans-Blot Turbo Transfer System (BioRad) following the manufacturers protocol. The membrane was blocked in 5% milk in TBS for 1 hour, and primary antibody (anti-TG2, Abcam 2386, 1:500 in 1.5% milk in TBS) overnight at 4° C. The blot was washed 3 x 15 minutes in TBS-T, and then incubated with secondary antibody (Goat anti-Mouse IgG (H+L) Cross-Adsorbed Secondary Antibody, Alexa Fluor™ 647, Invitrogen A-21235) at room temperature for 1 hour, followed by washing 3 x 15 minutes in TBS-T. The blot was imaged using a BioRad ChemiDoc MP, and then the blot was stained with amido-black stain to assess total protein loading.

#### Transamidation Activity Assay

100 ug of HeLa extract was diluted in low-salt buffer (20 mM HEPES pH 7.9, 150 mM KCl, 10 mM MgCl2), and 1X final transamidation assay buffer was added (25 mM Tris-HCl, pH 8, 10 mM CaCl2, 10 mM DTT, 10 mM KCl). Biotin-cadaverine (Millipore Sigma A5348) was added to a final concentration of 1 mM, and reactions incubated at 30° C for 2 hours. A western blot was run as described above, using a Streptavidin-Alexa Fluor™ 488 Conjugate (ThermoFisher S32354) to measure incorporation of biotin-cadaverine.

### Statistics

Statistical analyses for behavioral and immunoassay data were conducted using Prism software (GraphPad, v.10.4.1). Data distribution was assessed for normality. Data that met assumptions of normality were analyzed using parametric tests, while non-normally distributed data were analyzed using non-parametric alternatives. For experiments involving multiple conditions, one-way or two-way ANOVAs were performed, followed by *post hoc* analyses when appropriate. For time course analyses where multiple measurements were taken from the same animal, repeated measures ANOVAs were performed. Two-tailed Student’s t-tests were used for comparisons between two conditions. Behavioral data derived from manual observations and pregnancy/litter outcomes were analyzed using chi-square tests. Grubb’s test (alpha = 0.05) was applied to detect outliers where necessary. Statistical significance was defined as p ≤ 0.05.

### Bulk RNA-seq and analysis

#### RNA isolation and library preparation

Total mRNA was extracted from frozen brain tissues after homogenization in Trizol Reagent (Thermo Fisher) and cleaned using RNeasy Microcolumns (Qiagen) following the manufacturer’s instructions. For RNA-seq library preparation, 150 ng of mRNA per sample was used with either the Illumina Stranded mRNA Prep Kit (Illumina, #20040534) or the TruSeq RNA Library Prep Kit v2 (Illumina, #RS-122-2001), according to the manufacturer’s protocols. Library quality was assessed using a Qubit Fluorometer 2.0 (Thermo Fisher) and a High Sensitivity D5000 TapeStation assay (Agilent) before sequencing on a NovaSeq 6000 or NovaSeq X system.

#### Differential expression analysis

Raw fastq files, containing an average of 20–30 million reads per sample, were processed for pseudoalignment and abundance quantification using Kallisto (v. 0.46.1) against the Ensembl Mus musculus reference (v. 79)^79^. To filter lowly-expressed genes, only those with a total read count of at least 10 across all samples were retained. To account for unwanted variation among samples within each sequencing experiment that could arise from technical or biological factors unrelated to the conditions of interest (including litter size, estrous stage, day of sample collection, etc.), RUVs (v1.32.0) was applied with a negative control gene set derived from the total genes identified per sequencing experiment, after ensuring that unwanted variation did not correlate with covariates of interest, as described previously^80,81^. Differential expression analysis was performed using DESeq2 (v1.38.3)^82^, with significant genes defined by an adjusted p-value < 0.05. For brain-wide transcriptome comparisons in which samples were processed across multiple sequencing runs and stemming from separate cohorts, pairwise comparisons were performed independently for each brain region. For all other experiments, where subjects came from the same cohort and were processed in a single sequencing run, all groups were analyzed together to maintain consistent normalization within the experiment. Gene expression time course analyses examining the periods before, during, and after pregnancy and postpartum were performed on normalized count data using the ImpulseDE2 package (v0.99.10)^31^ for each brain region. Significant genes exhibiting transient regulation or monotonous changes in expression were identified using case-only differential expression analysis, with a Q-value threshold of 0.05.

#### Weighted gene co-expression analysis

To identify brain-wide gene co-expression networks, normalized count data for all brain regions were compiled and analyzed using the WGCNA package (v1.73)^20^. Co-expression networks were constructed from the 7,500 most variable genes, determined by ranking gene variance. A soft threshold power of 12 was identified with the pickSoftThreshold function to ensure scale-free network properties. Modules were identified based on dissimilarity of a signed topological overlap matrix (TOM), and named with an arbitrary color. The “gray” module encompassed genes that did not segregate into any specific module, and was therefore removed from further analyses. To assess the enrichment of differentially expressed genes (DEGs) within each brain region per module, Fisher’s exact tests were performed using the fisher.test() function in R (v4.3.0). To analyze the correlation between gene modules and brain regional sensitivity to parity, brain regions were categorized into two groups based on the top five regions with significant DEG overlap. These regions were designated as “High Sensitivity” regions, while all other regions were classified as “Low Sensitivity.” Pearson correlation coefficients were calculated to examine the relationship between each module and the trait (regional sensitivity × parity status), with statistical significance determined by Student’s t-tests (p <0.05). Heatmaps were generated to visualize these module-trait correlations, and module-specific gene lists were exported for pathway enrichment analysis.

#### Pathway and predicted upstream regulator analyses

Functional annotation of DEGs was conducted using ShinyGO (v0.81)^83^, with all protein-coding genes in the mm10 genome used as the background. Pathways with an FDR < 0.05 were considered significant. All significant pathways and associated statistics are provided in the Supplementary Tables. For figures, relevant pathways were selected from the top 10 or 1/3 of significant terms, ranked by FDR, to emphasize processes consistent with hypotheses informed by published literature. For GO term selection, Revigo^84^ was used to reduce redundancy of overlapping GO terms when needed. Ingenuity Pathway Analysis (IPA; Qiagen, Inc., v.01-23-01) was used to predict upstream regulators for DEG lists^85^. For “High Sensitivity” and “Low Sensitivity” regions (see *WGCNA* section), DEGs shared by at least two or three brain regions were extracted, irrespective of the direction of change. The IPA software was used to identify upstream regulators associated with each DEG list, with statistical significance defined as p < 0.05. To investigate the mechanisms underlying parity programming of dHpF plasticity, significant upstream regulators were systematically prioritized based on one or more of the following criteria: (1) multiple molecules involved in the same signaling pathways (e.g., progesterone and its receptor, PGR); (2) structurally similar molecules that engage analogous signaling cascades (e.g., levodopa and dopamine); (3) molecules known to be influenced by reproductive exposures; (4) molecules demonstrated to play a role in hippocampal plasticity; (5) molecules identified as significant in both “High Sensitivity” and “Low Sensitivity” regulator lists; and (6) molecules with high statistical significance values. Following selection, upstream regulators were categorized into general molecular classes (e.g., “Hormone,” “Transcription Factor,” “Lipid”) and grouped for visualization. Data are presented as a bubble plot, with marker shape representing number of brain regions sharing DEGs and marker size corresponding to significance.

#### Gene expression overlap analyses

Jaccard indices were calculated for overlapping gene lists using the GeneOverlap package (v1.36.0)^86^, with significance defined by *p* < 0.05. Fisher’s exact tests were calculated using the base *fisher.test()* function in R (v. 4.3.0) following construction of contingency tables for each comparison. Transcriptome-wide, threshold-free gene expression overlap was visualized using Rank-Rank Hypergeometric Overlap (RRHO) heatmaps generated with the RRHO2 package (v1.0)^30^. Gene lists were ranked by signed p-values, calculated as the log10-transformed nominal p-value multiplied by the sign of the fold change, without applying differential expression thresholds.

#### Cell-type deconvolution

Brain cell-type proportion was estimated from normalized expression data using the BRETIGEA package (v.1.0.4)^18^ with default markers. For comparisons between groups, surrogate proportion variables were normalized using the Normalize function in Prism (GraphPad, v. 10.4.1). Each sub column was normalized separately, with the smallest value set to 0% and the largest value set to 100%. To calculate log_2_(fold change) between groups, normalized values per cell-type per group were averaged and expressed as the log2 ratio of the group means.

### Single nuclei RNA-seq

#### Nuclei isolation and library preparation

For each animal, 2 mm dHpF-enriched tissue micropunches were collected bilaterally from consecutive 1 mm slices from -0.80 to -2.80 mm relative to bregma for a total of 4 punches (Ted Pella). Samples were processed in batches of 4-6, with each group represented within each batch. Nuclei were isolated using a modified version of a sucrose density gradient isolation protocol^87^. Briefly, thawed tissues were placed in 1 mL of lysis buffer (0.32 M sucrose, 5 mM CaCl_2_, 3 mM magnesium acetate, 0.1 mM EDTA, 10 mM Tris-HCl pH 8, 1 mM DTT, 0.1% Triton X-100) with 50 μL of 25 U/mL RNase inhibitor (Takara #2313B) in a dounce homogenizer (Wheaton #357538). Homogenization was performed with 20 strokes using a tight pestle. Another 1 mL of lysis buffer was added, followed by an additional 10 strokes. The resulting 2 mL homogenate was transferred to a 15 mL Open-Top Thinwall Polypropylene Tube (Beckman-Coulter #361707). The homogenizer and pestle were rinsed with 2 mL of lysis buffer, and this wash was combined with the homogenate for a total of 4 mL. The homogenate was carefully underlaid with 9 mL of sucrose solution (1.8 M sucrose, 3 mM magnesium acetate, 1 mM DTT, 10 mM Tris-HCl pH 8) and ultracentrifuged at 24,000 rpm for 1 hour at 4 °C using a Sorvall™ WX+ centrifuge. After centrifugation, the supernatant and debris at the interface were gently removed. The nuclear pellet was resuspended in 1 mL of resuspension buffer (0.02% bovine serum albumin in DPBS with 25 μL of 25 U/mL RNase inhibitor) and incubated on ice for 10 minutes. The suspension was passed through a 35 μm nylon mesh filter (Corning #352235) into a 1.5 mL RNase/DNase-free microcentrifuge tube and centrifuged at 2600 x g for 10 minutes at 4 °C. Supernatants were discarded, and nuclei were resuspended in 200 μL of resuspension buffer. A 10 μL aliquot of the nuclei suspension was stained with Trypan Blue to assess quality and concentration using a Countess 3 Automated Cell Counter. Nuclei suspensions were loaded onto a Chromium Single Cell 3′ chip (10X Genomics, v3) and processed according to the manufacturer’s protocol, targeting 10,000 nuclei. Single-nuclei libraries were generated using the 10X Chromium Next GEM Single Cell 3′ v3.1 (Dual Index) protocol (CG000315 Rev A). Libraries were pooled, loaded onto a single 10B 100 Cycle Flowcell, and sequenced using an Illumina NovaSeq 6000 system to generate 25-30,000 paired-end 2 × 100 bp reads.

#### Data analysis

FastQ files were processed with the 10X Genomics Cell Ranger pipeline (v7.1.0) to demultiplex reads, align them to the mouse genome (mm10-2020-A), remove PCR duplicates, and generate gene expression matrices. Cell Ranger filtered outputs were analyzed using Seurat v4.3.0^88^, and mitochondrial RNA content per cell was calculated using the GRCm39 (mm10) genome annotation and regressed out using SCTransform normalization protocol included in the Seurat toolkit with 20 principal components (PCs) and a resolution of 0.1. To estimate ambient RNA and correct for background contamination, the SoupX (v1.6.2) package^89^ was used for each sample using raw and filtered feature matrices from the Cell Ranger output. Heterotypic doublets were identified and removed using DoubletFinder (v2)^90^ to ensure the integrity of singlet datasets. Filtered singlet datasets were then re-normalized and integrated using the same Seurat SCTransform v2 workflow mentioned above. Cell clusters were annotated using a combination of expert curation based on published marker genes^37,91–96^, and label transfer from hippocampal reference datasets, including the Allen Brain Map and Broad Institute resources^36,37^. Clusters with contaminant cell populations expressing markers for choroid plexus (*Ttr*), ependymal (*Tmem212*), and vascular leptomeningeal cells (*Vtn, Col1a2*) were removed from the analysis^96–99^. Additionally, as the sequential 2mm micropunches encompassed portions of cortical, thalamic and vHpF regions, clusters characterized by enrichment of published non-dHpF neuronal markers using Seurat’s FindMarkers function (layer 5/6 cortical: *Rorb, Foxp2*^99^; ventral granule neurons: *Tox3*^93^) were also removed from the analysis. Cell cluster proportion analyses were conducted using the scProportionTest package^100^, which employs a Monte Carlo permutation test to evaluate whether observed differences result from random sampling. Proportional differences between conditions were compared to a null distribution generated by resampling, and statistical significance was determined by permutation-based p-values with confidence intervals estimated via bootstrapping. Differential expression analysis was conducted using pseudobulk analysis, where gene counts were summed across all cells within each sample for each cell type cluster using the AggregateExpression() function. DESeq2 was then applied at the sample level to conduct differential expression. To explore pathways underlying cluster-specific differences across conditions, pathway analysis was conducted using ShinyGO on genes meeting the following criteria: log2FC > 1.5 and p < 0.05.

### CUT&RUN-seq

#### Cleavage Under Targets and Release Using Nuclease

The procedure was adapted from established protocols^46,101^. For mouse brain tissue, two unpooled 1.5mm dHpF punches were used for each biological replicate and split across three reactions (H3K4me3Q5ser, H3K4me3, IgG). For human brain tissue, ∼10mg of tissue dissected from individual frozen subiculum samples was collected from each subject, and split across the indicated antibodies. Samples were dounce homogenized in nuclear extract (NE) buffer (20mM HEPES-KOH pH 7.9, 10mM KCl, 0.5mM spermidine, 0.1% Triton-X, 20% glycerol with protease inhibitors) and passed through a 21 gauge needle 10x. Nuclei were pelleted at 1,100g for 5 min at 4 °C in a swinging-bucket rotor, passed through a 40 μM strainer (pluriSelect USA), washed again in 500 μl NE buffer and counted. BioMag Plus Concanavalin A beads (Polysciences) were prepared per reaction by washing three times with binding buffer (20 mM HEPES-KOH pH 7.9, 10 mM KCl, 1 mM CaCl₂, 1 mM MnCl₂). Beads (15 μl) were aliquoted into 1.7 ml DNA low-bind tubes (Eppendorf) containing 500 μl NE buffer and 100,000 nuclei per reaction. Samples were rotated at room temperature for 10 min, then bead-bound nuclei were washed three times with wash buffer (20 mM HEPES pH 7.5, 150 mM NaCl, 0.1% Triton X-100, 0.1% Tween-20, 0.5 mM spermidine, 0.1% BSA, and protease inhibitors), resuspended in 100 μl antibody buffer (wash buffer with 2 mM EDTA), and mixed. 2 μl antibodies were added to the corresponding tubes: H3K4me3 (Active Motif, 39159), H3K4me3Q5dopaminyl (Millipore, ABE2590), or rabbit IgG (Invitrogen, 10500c). Samples were incubated overnight at 4 °C on a rotating mixer angled upward at 20 degrees. The next day, nuclei were washed twice with cold wash buffer, and incubated with 2.5 μl pAG-MNase (Epicypher, 15-1016) for 1 h at 4 °C. After four washes with cold wash buffer and one with low-salt rinse buffer (20 mM HEPES pH 7.5, 0.5 mM spermidine, 0.1% Tween-20, 0.1% Triton X-100), nuclei were resuspended in calcium incubation buffer (3.5 mM HEPES pH 7.5, 10 mM CaCl₂, 0.1% Tween-20, 0.1% Triton X-100) and placed into an ice-cold block at 4 °C. MNase digestion was stopped by adding 100 μl of 2x Stop Buffer (340 mM NaCl, 20 mM EDTA, 5 mM EGTA, 0.1% Tween-20, 0.1% Triton X-100, 25 μg/ml RNase A, and 0.05 ng/100 μl E. coli spike-in DNA), followed by a 15 min incubation at 37 °C without shaking. Beads were then placed on a magnet and 200 μl of supernatant was collected. DNA was purified using the Zymo ChIP DNA Clean & Concentrator kit (D5205), eluted in 30 μl, and stored at −20 °C for library preparation. Libraries were generated using the NEBNext Ultra II DNA library kit, quantified using a Qubit fluorometer with the HS DNA kit, checked for size distribution on the Agilent TapeStation, pooled equimolarly, and sequenced on an Illumina NovaSeq X.

#### Data analysis

Raw fastq files were aligned to the hg19 or mm10 genome using bowtie2 (v2.5.0)^102^. Low-quality reads were filtered using Samtools (v.1.9) with a MAPQ cut-off score of 30^103^. Only unique, deduplicated reads were retained for further processing. Bigwig files were produced using the deepTools package (v.3.5.1), using an ENCODE hg19 or mm10 v2 blacklist file to discard regions with consistently non-specific signal, and scaled using *E. coli* spike-in controls to normalize sequencing depth. To determine normalization factors based on *E. coli* reads, each sample was aligned to the *E. coli* genome (MG1655), and the unique deduplicated reads were compared across groups per antibody per experiment. The sample with the lowest number of *E. coli* reads was determined (“minimum”), and all samples were scaled by dividing their corresponding *E. coli* read count by this minimum number^104^. For each group, bigwig files were merged and peak callling was conducted using MACS2 (v2.1.0) with the corresponding merged IgG file as control, filtered for peaks with FDR < 0.05^105^. Peak annotation was conducted using HOMER (v4.1.1)^106^. Heatmaps were made either using the DiffBind (v3.8.4) or deepTools (v3.5.5) packages^53,107^. For deepTools, heatmaps were made by merging DEGs from RNA-seq data with TSSs downloaded from the UCSC genome browser using the canonically annotated transcript for each gene. Profiles were generated and statistically analyzed using the deepStats package^108^ by using the dsCompareCurves function to perform Wilcoxon Rank-sum tests per-bin. For DiffBind analysis, heatmaps were made for peaks identified by DiffBind’s differential peak algorithm, where differential peaks were first filtered using a log_2_(fold change) threshold > 0.1 and defined at p < 0.05, where log_2_(fold change) was calculated as log2(parity) − log2(NP), based on prior empirical observations used to define thresholds for differential peaks^49^. ChEA analysis on annotated loci was conducted using EnrichR with a significance threshold of adjusted *p* < 0.05^109^.

## DATA AND MATERIALS AVAILABILITY

The genomics data generated in this study have been deposited in the National Center for Biotechnology Information Gene Expression Omnibus (GEO) database. We declare that the data supporting findings for this study are available within the article and Supplementary Information. Related data are available from the corresponding author upon reasonable request. No restrictions on data availability apply.

---

**Extended Data 1:**
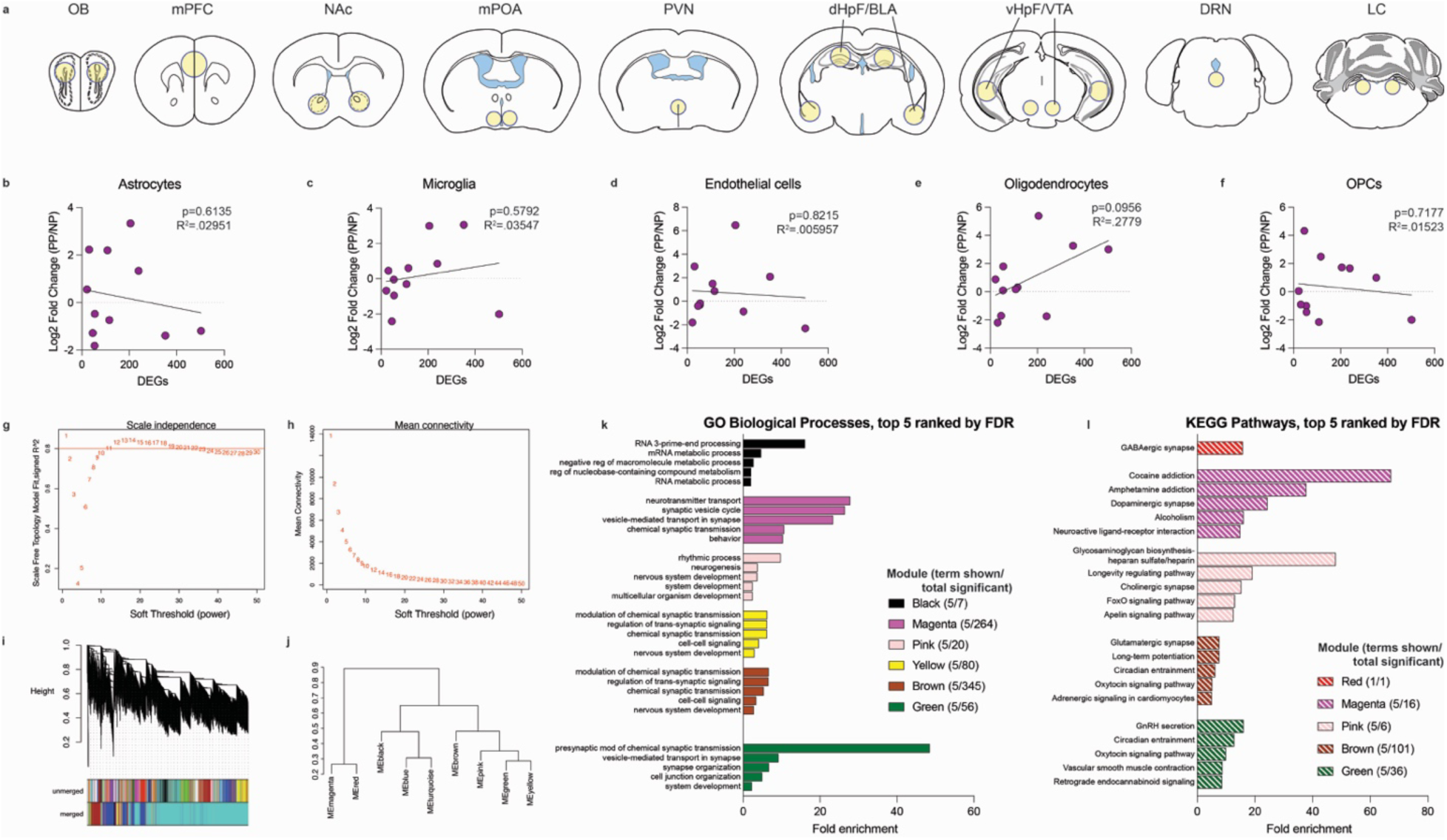
Brain-wide transcriptomic analysis of cell markers and gene expression networks. **a)** Brain regions selected for bulk transcriptional profiling: olfactory bulb (OB), medial prefrontal cortex (mPFC), nucleus accumbens (NAc), medial preoptic area (mPOA), paraventricular nucleus (PVN), dorsal hippocampal formation (dHF), basolateral amygdala (BLA), ventral hippocampus (vHF), ventral tegmental area (VTA), dorsal raphe nucleus (DRN), and locus coeruleus (LC). Yellow circles indicate area selected for tissue micropunching. **B-F)** Nonsignificant correlations between the fold change (NP *vs.* PP) in normalized surrogate proportion variables generated from cell-type deconvolution of bulk RNA-seq data for **B)** astrocytic, **c)** microglia, **d)** endothelial, **e)** mature oligodendrocyte, and **f)** oligodendrocyte precursor cell markers with the number of DEGs identified from each brain region. **g)** Scale independence plot for soft-threshold power selection for WGCNA analysis, illustrating the relationship between β and the scale-free topology fit index, identifying the optimal β value = 12. **h)** Mean connectivity plot illustrating the average network density for each evaluated soft-thresholding power. **i** Dendrogram of gene clustering based on topological overlap, highlighting the hierarchical clustering process used for module detection and merging of similar modules. **j)** Dendrogram of eigengene network adjacency depicting similarity of modules. **k, l)** Significantly enriched **k)** GO Biological Processes and **l)** KEGG pathways, sorted by FDR values, from module gene sets. Bars are color-coded to indicate modules. The total number of significant terms for each analysis is provided.

**Extended Data 2:**
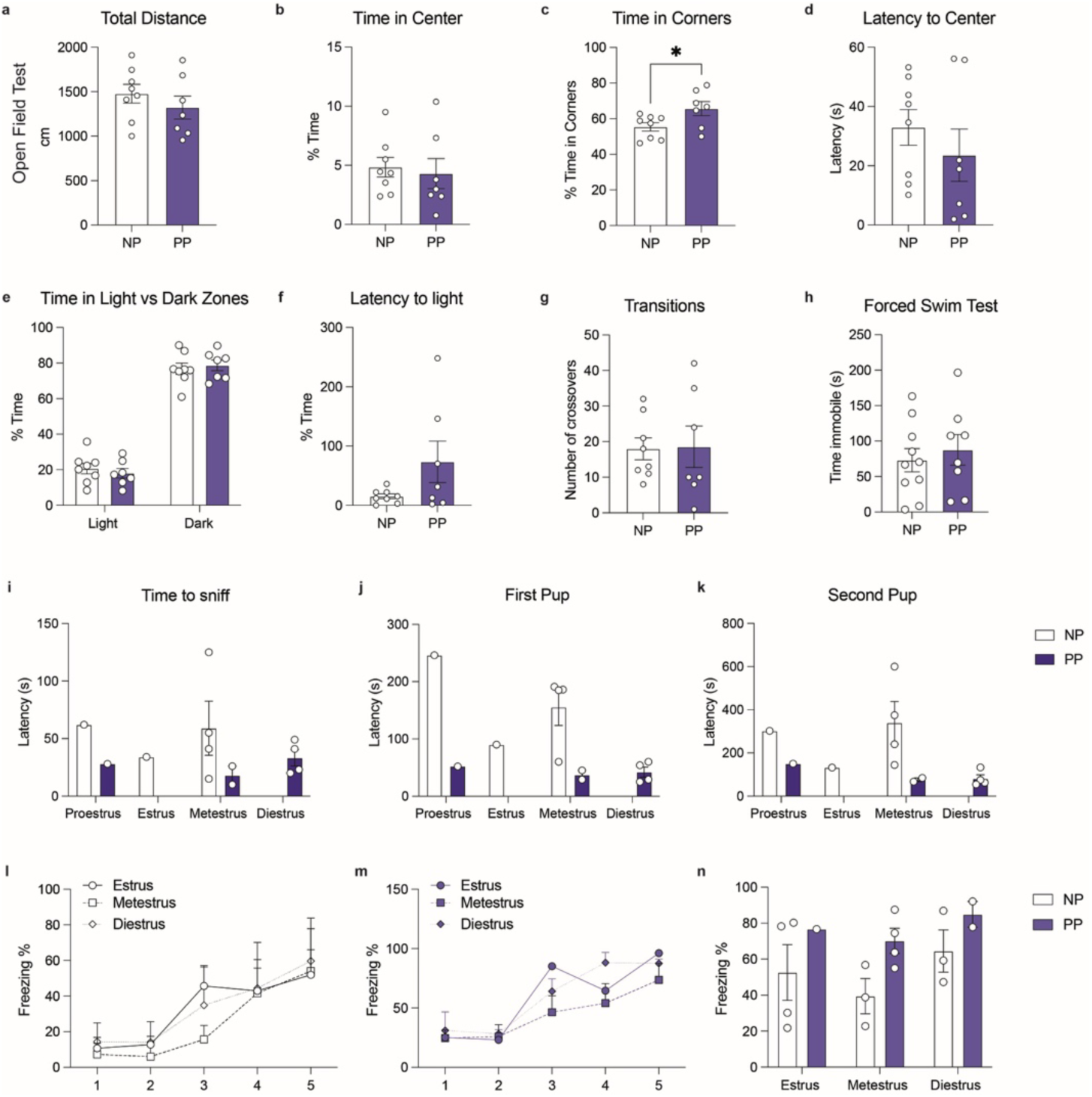
Parity associated behavioral adaptations are not influenced by locomotion, anxiety-/depressive-like behaviors, or estrous stage. a-d) In the open field test, there was no change in total distance, time spent in the center, or latency to the center. While there was a significant increase in time spent in corners for PP dams (Student’s t-test, t(13) = 2.307, *p = 0.0382), this did not impact any other outcomes on the open field test. **e-g)** In the light-dark box, there was no difference in time spent in lights *vs.* dark zones, latency to enter the light zone, or transitions between zones. **h)** There was no difference in time spent immobile on the forced swim test. **i-k)** Stratification of pup retrieval behaviors by estrous stage on the day of testing did not reveal a significant effect of estrous stage. **l-n)** Similarly, stratification of contextual fear conditioning outcomes did not identify a significant effect of estrous stage on acquisition or context recall in NP or PP groups. Error bars represent mean ± SEM. N=6-11 animals/group.

**Extended Data 3:**
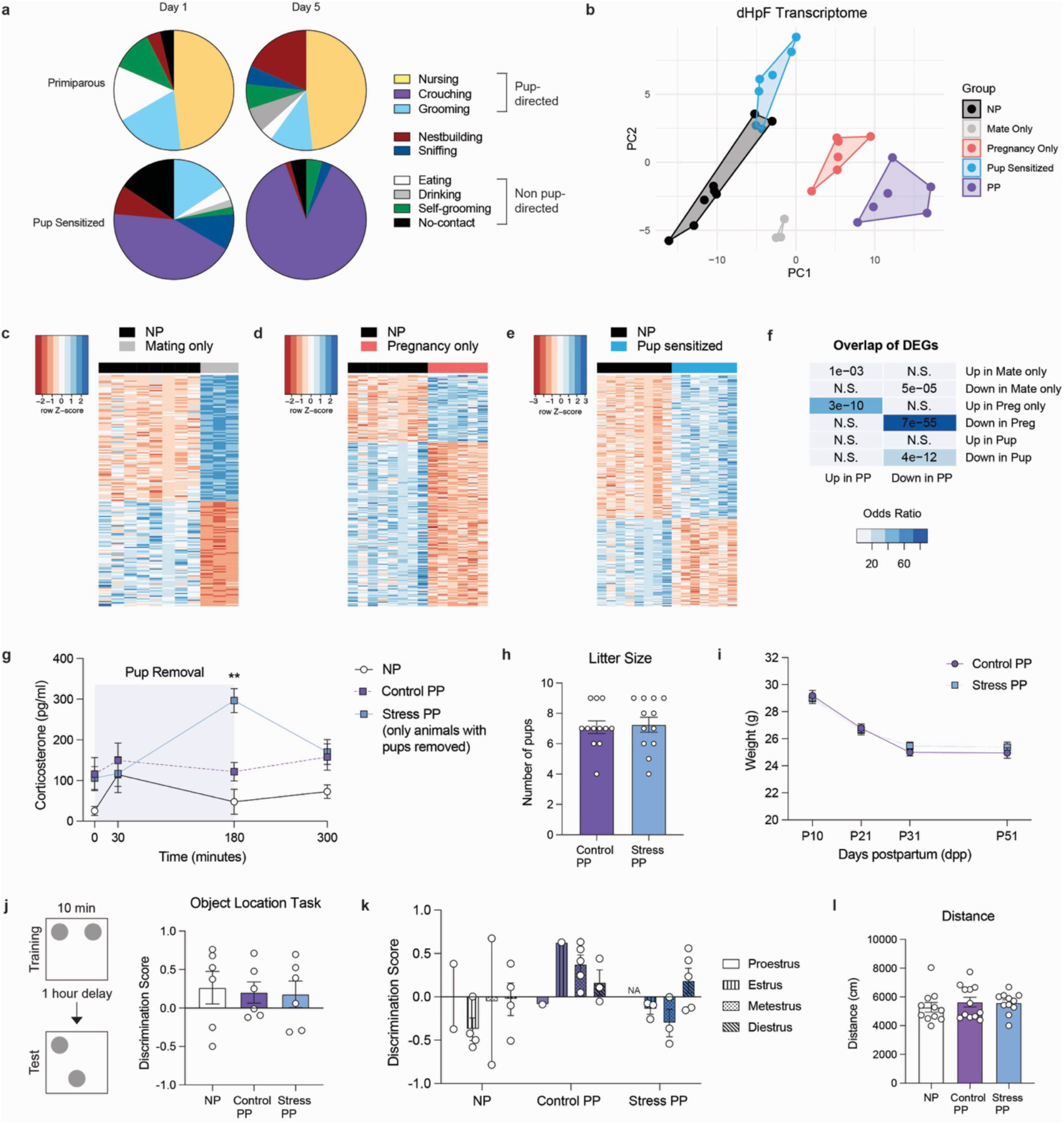
Postpartum experiences modulate the extent of parity-associated dHF transcriptomic and behavioral adaptations. **a)** Pie charts showing significant shifts towards pup-directed behaviors, representing maternal behavior initiation after four days of pup exposure in Pup Sensitized virgin females (χ²(7) = 33.52, p = 2.11×10^-5^). **b)** Principle components analysis of NP *vs.* PP DEGs (adj p < 0.05) from dHF, showing the Pregnancy Only group clusters most closely with PP. **c)** Heatmaps of all significant DEGs comparing NP *vs.* Mating only, **d)** NP *vs.* Pregnancy only, **e)** NP *vs.* Pup sensitized dHF transcriptomes. **f)** Odds ratio analysis of DEG overlap for all comparisons (*vs.* NP). Insert numbers indicate respective *p* values for each association (N.S., *p* > 0.05). **g)** Pup separation increases maternal corticosterone levels over 3-hours (RM two-way ANOVA, group: F(2,15) = 12.86, p = 0.0006, time: F(2.125, 31.87) = 2.759, p = 0.0756, interaction: F(6,45) = 3.294, p = 0.009; Tukey’s multiple comparisons test: Control *vs.* Stress PP, **p < 0.01). N = 6 animals/group. **h)** No effect of postpartum stress on litter size. N = 12/group. **i)** Postpartum stress did not alter maternal weights. **j)** All groups discriminated the novel location following 10-minutes of training on the object location task (one way ANOVA, F(2,15) = 0.06452, p = 0.9378) N=6 animals/group. **k)** Stratification of object location test scores did not show a significant effect of estrous stage. **l)** There was no difference in the total distance travelled in the open field test across groups. Error bars represent mean ± SEM.

**Extended Data 4:**
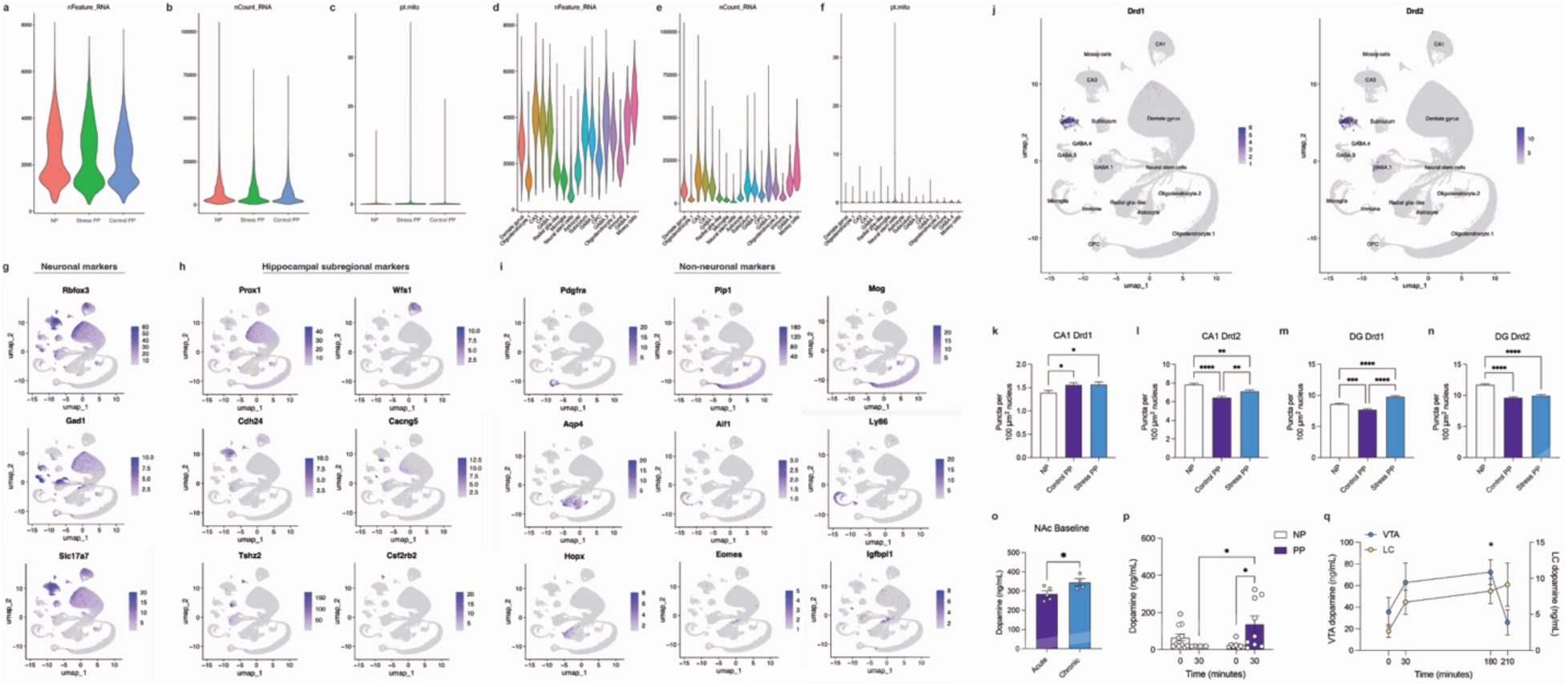
snRNA-seq quality control assessments, cell marker gene expression, and postpartum dopamine modulation. Violin plots showing the **(a)** number of detected genes, **(b)** UMI number, and **(c)** mitochondrial gene % by group following regression for mitochondrial gene mapping. (**d-f)** Breakdown of these metrics by annotated clusters show <10% mitochondrial gene across all cell-types, except for a minority of neural stem cells. Unsupervised clustering of snRNA-seq data with cells colored for **(g)** neuronal, excitatory, and inhibitory markers; **(h)** hippocampal subregional markers; and **(i)** non-neuronal markers including for glial and neural progenitor populations. **j)** Unsupervised clustering of snRNA-seq data with cells colored for *Drd1* (left) and *Drd2* (right) expression. **k)** Quantification of *Drd1* mRNA puncta in CA1 nuclei (one-way ANOVA, (F(2,19204) = 3.963, p = 0.019). **l)** Quantification of *Drd2* mRNA puncta in CA1 nuclei (one-way ANOVA, (F(2,19204) = 23.16, p < 0.0001). **m)** Quantification of *Drd1* mRNA puncta in dentate gyrus nuclei (one-way ANOVA, (F(2,21864) = 42.09, p < 0.0001). **n)** Quantification of *Drd2* mRNA puncta in dentate gyrus nuclei (one-way ANOVA, (2,21864) = 41.79, p < 0.0001). Tukey’s multiple comparisons test, *p < 0.05, **p < 0.01, ***p < 0.001, ****p < 0.0001. **o)** Chronic postpartum stress resulted in higher baseline levels of dopamine in the nucleus accumbens (NAc; Student’s t-test, t(7) = 2.568, *p = 0.0371). N= 4-5 animals/group. **p)** Dopamine response is not induced in NP dHF 30-minutes following littermate separation (two-way ANOVA, interaction of time x group (F(1,30) = 8.78, p = 0.0059, time (F(1,30) = 1.47, p = 0.2359, group, (F(1,30) = 2.054, p = 0.1621); Tukey’s multiple comparisons test, *p < 0.05). N = 6-11 animals/group. **q)** Dopamine levels increased during pup separation in the VTA (left y-axis), but not in the locus coeruleus (right y-axis), resembling patterns of NAc and dHF dopamine dynamics (two-way ANOVA, effect of brain region (F(1,46) = 32.32, p < 0.0001), time (F(3,46) = 2.311, p = 0.0866), interaction (F(3,46) = 1.118, p = 0.1109); Dunnett’s multiple comparisons test, *p < 0.05). N = 5-8 samples/group. Error bars represent mean ± SEM.

**Extended Data 5:**
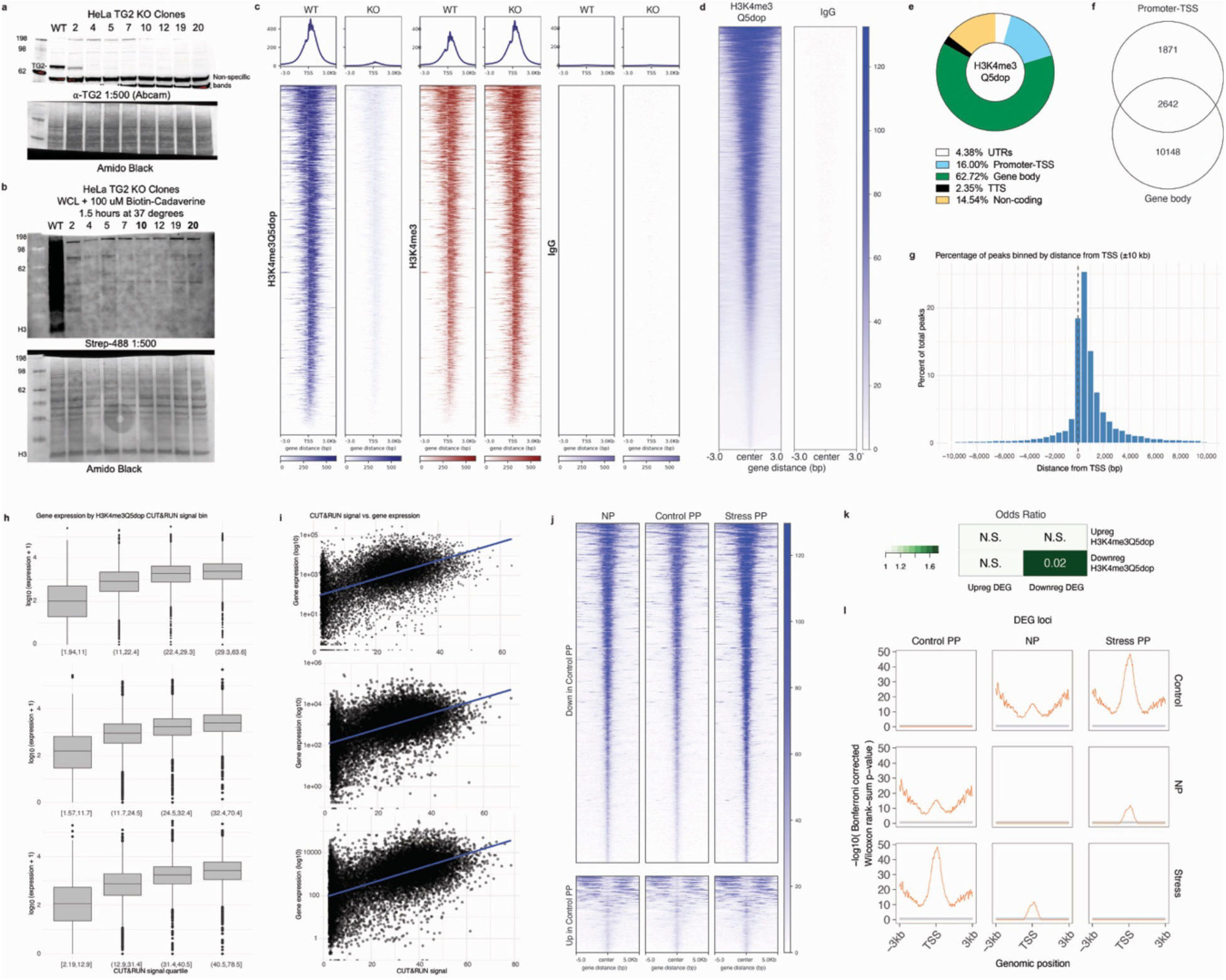
H3K4me3Q5dop enrichment corresponds with gene expression. a,b) Validation of TG2 knockout (KO) cell lines using **(a)** Western blot analysis to confirm the absence of TG2 protein, and **(b)** a transamidation activity assay to show loss of biotin-cadaverine incorporation in TG2 KO cells, indicating loss of enzymatic activity. **c)** TG2 KO results in a selective >90% reduction of H3K4me3Q5dop enrichment (*vs.* H3K4me3 alone) at transcription start sites (TSS) compared to WT controls. **d)** Heatmaps of H3K4me3Q5dop and IgG peak enrichment at all enriched loci in NP dHF tissue. **e)** Distribution of H3K4me3Q5dop enrichment occurs mainly at genic loci (83%), with **f)** overlap occurring between uniquely annotated promoter-TSS and gene body (exon/intron) regions and **g)** ∼63.4% of total signal occurring within 2kB of the TSS. **h)** Boxplots displaying log₁₀-transformed gene expression across quartiles of H3K4me3Q5dop signal for NP (top), Control PP (middle), and Stress PP (bottom). Significant differences were observed across bins (Kruskal–Wallis *p* < 2.2 × 10⁻¹⁶) for each group between each quartile. Boxes represent the interquartile range with the median indicated, and whiskers show the rangs with outliers plotted as individual points. **i)** Scatter plots showing the correlation between H3K4me3Q5dop CUT&RUN signal (maximum per gene) and mean gene expression (log₁₀-transformed) for NP (top), Control PP (middle), and Stress PP (bottom). Significant positive correlations were observed for each group (NP: Spearman’s ρ = 0.55, *p* < 2.2 × 10⁻¹⁶; Control PP: Spearman’s ρ = 0.52, *p* < 2.2 × 10⁻¹⁶; Stress PP: Spearman’s ρ = 0.56, *p* < 2.2 × 10⁻¹⁶), indicating that increased H3K4me3Q5dop enrichment corresponds with higher transcriptional output. **j)** Heatmaps of all differential peaks (*p* < 0.05; log_2_FoldChange ≥ |0.1|) between NP/Stress PP vs., separated by directionality and centered on genomic regions to show the majority of altered peaks decrease in Control PP dHF. **k)** Odds ratio analysis of differential H3K4me3Q5dop peaks and differentially expressed genes show significant association between downregulated histone dopaminylation and gene expression changes. Insert numbers indicate respective p values for each association (N.S., *p* > 0.05). **l)** Statistical comparison of H3K4me3Q5dop profiles across groups at loci associated with DEGs. Each panel shows the –log₁₀ Bonferroni-corrected *p*-values from bin-wise Wilcoxon rank-sum tests comparing CUT&RUN signal across ±3 kb surrounding TSSs of DEG-associated loci. Comparisons are shown for each pairwise group contrast. Gray bars indicate bins that did not reach statistical significance after multiple hypothesis correction. The orange line reflects the statistical significance of signal differences across the TSS window, with higher values indicating greater confidence in differential enrichment between groups.

**Extended Data 6:**
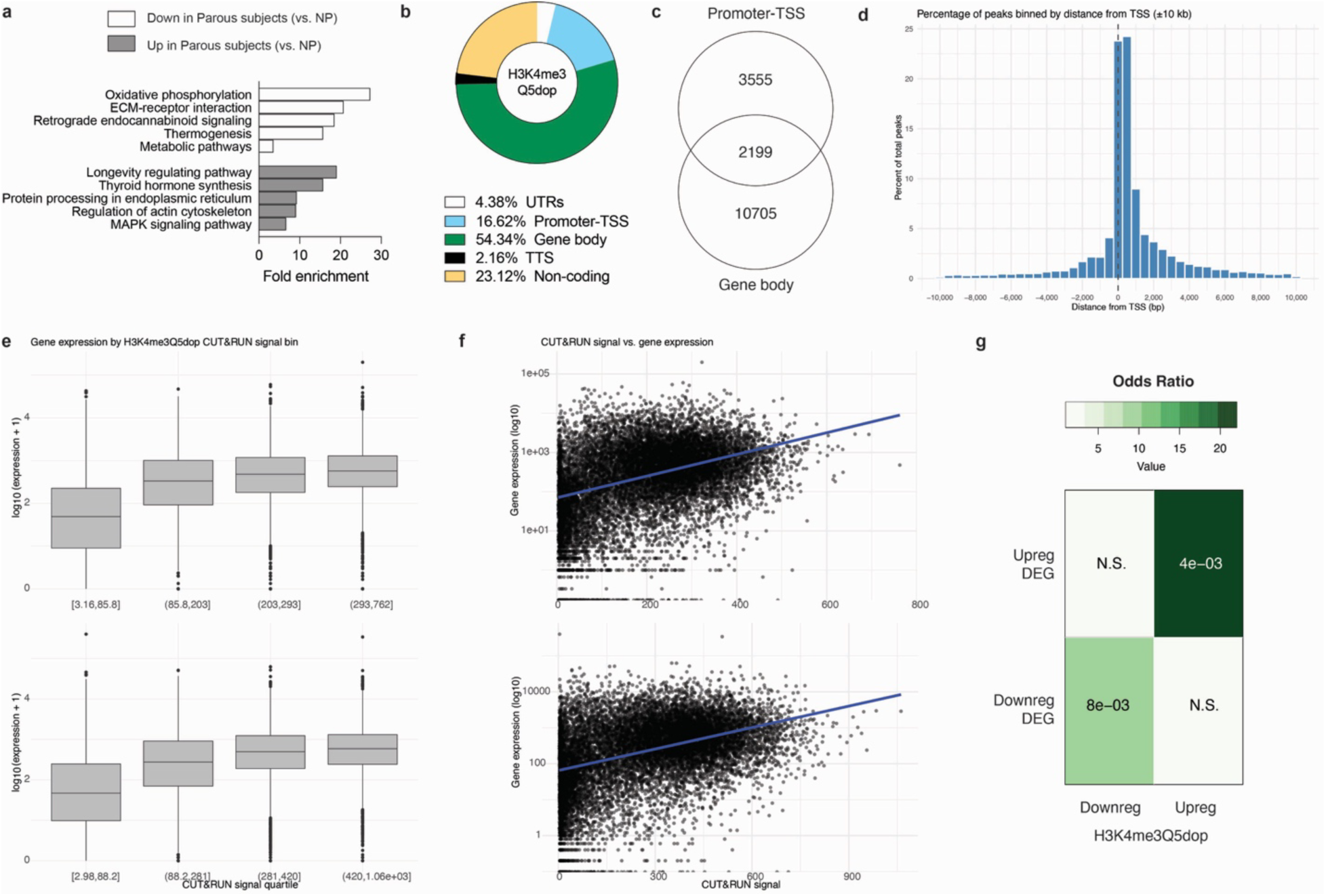
H3K4me3Q5dop enrichment in human brain. **a)** Pathway analysis for DEGs of NP vs. parous human dSub tissues by directionality (padj < 0.05). **b)** Distribution of H3K4me3Q5dop enrichment occurs mainly at genic loci (75.3%), with **c)** overlap occurring between uniquely annotated promoter-TSS and gene body (exon/intron) regions, similar to in mouse dHF, and **d)** ∼52.0% of total signal occurring within 2kB of the TSS. **e)** Boxplots displaying log₁₀-transformed gene expression across quartiles of H3K4me3Q5dop signal for NP (top) and parous subjects (bottom). Significant differences were observed across bins (Kruskal–Wallis *p* < 2.2 × 10⁻¹⁶) for each group and between each quartile. Boxes represent the interquartile range with the median indicated, and whiskers show the rangs with outliers plotted as individual points. **f)** Scatter plots showing the correlation between H3K4me3Q5dop CUT&RUN signal (maximum per gene) and mean gene expression (log₁₀-transformed) for NP (top) and parous subjects (bottom). Significant positive correlations were observed for each group (NP: Spearman’s ρ = 0.41, *p* < 2.2 × 10⁻¹⁶; Parous: Spearman’s ρ = 0.40, *p* < 2.2 × 10⁻¹⁶), indicating that increased H3K4me3Q5dop enrichment corresponds with higher transcriptional output. **g)** Odds ratio analysis of differential H3K4me3Q5dop peaks and differentially expressed genes show significant association between changes in H3K4me3Q5dop signal and gene expression changes. Insert numbers indicate respective p values for each association (N.S., p > 0.05).

**Extended Data 7:**
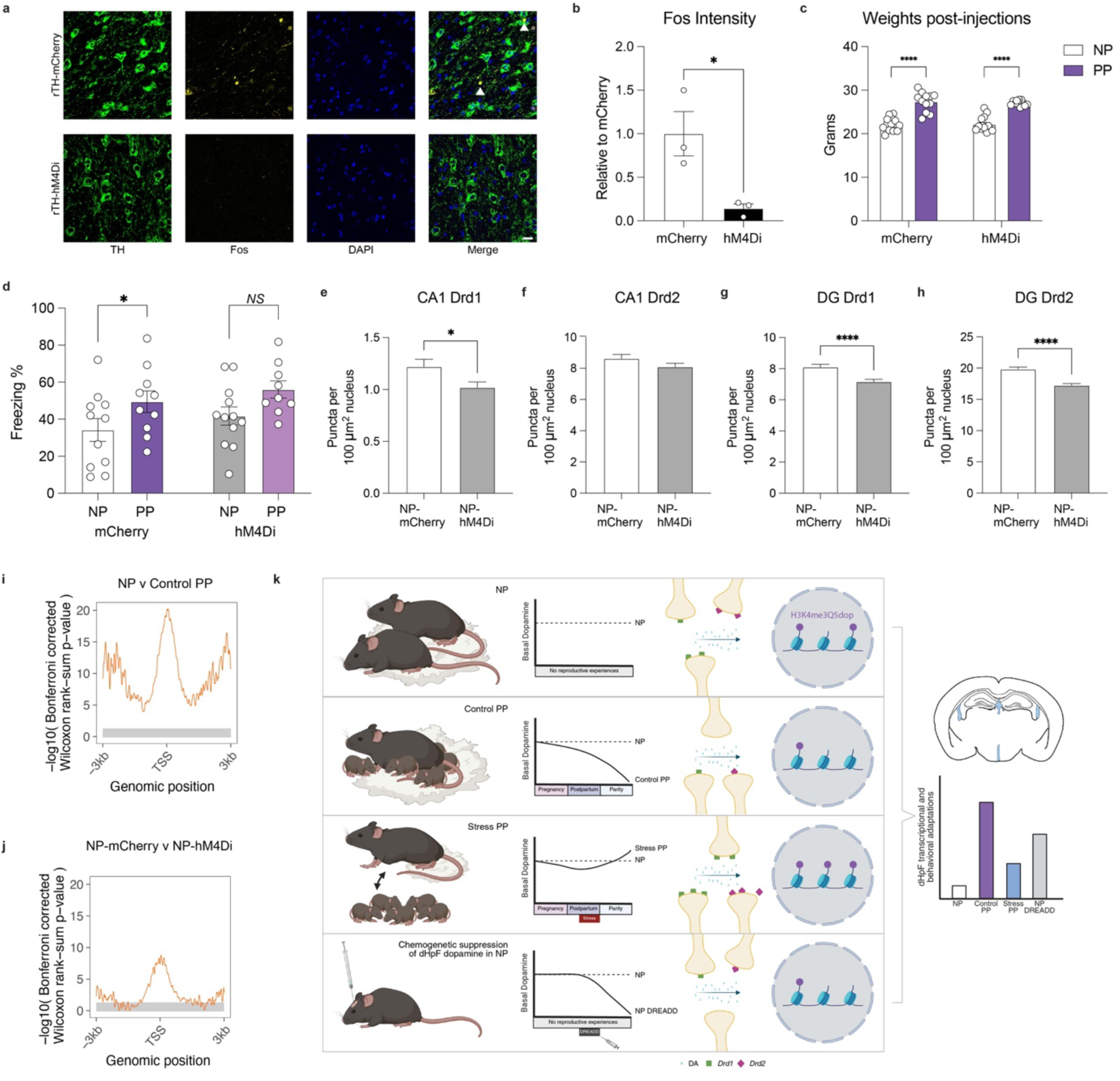
Inhibition of VTA-dHF projection mimics sustained changes in dopamine receptor dynamics. **a)** Representative images of c-Fos immunostaining in the VTA. Scale bars, 20 μm. **b)** Quantification of c-Fos immunoreactivity in VTA TH+ neurons (Student’s t-test; t(4) = 3.299, *p = 0.03). N = 3 animals/group. **c)** Chronic DCZ injections did not alter weights at 21 dpp (two-way ANOVA; main effect of group (F(1,48) = 128, p < 0.0001), virus (F(1,48) = 0.0862, p = 0.7702); Tukey’s multiple comparison’s test, ****p < 0.0001). **d)** Parity induced significantly greater freezing on the context recall text in mCherry dams, with trending effects between hM4Di groups (two-way ANOVA (effect of group, (F(1,38) = 7.177, p = 0.0109, virus (F(1,38) = 1.631, p = 0.2093, Fisher’s LSD, *p £ 0.05). N = 9-12/group. **e)** Quantification of *Drd1* mRNA puncta in CA1 nuclei (Student’s t-test, (t(4854) = 2.249, *p = 0.00246). **f)** Quantification of *Drd2* mRNA puncta in CA1 nuclei (Student’s t-test, (t(4854) = 1.502, p = 0.1132). **g)** Quantification of *Drd1* mRNA puncta in dentate gyrus nuclei (Student’s t-test, (t(11946) = 3.900, ****p < 0.0001). **h)** Quantification of *Drd2* mRNA puncta in dentate gyrus nuclei (Student’s t-test, (t(11946) = 6.034, ****p < 0.0001). **i, j)** Statistical comparison of H3K4me3Q5dop profiles between NP-mCherry and NP-hM4Di at loci associated with **(i)** NP *vs.* Control PP DEGs and **(j)** NP-mCherry *vs.* NP-hM4Di DEGs. Each panel shows the –log₁₀ Bonferroni-corrected *p*-values from bin-wise Wilcoxon rank-sum tests comparing CUT&RUN signal across ±3 kb surrounding TSSs of DEG-associated loci. Comparisons are shown for each pairwise group contrast. Gray bars indicate bins that did not reach statistical significance after multiple hypothesis correction. The orange line reflects the statistical significance of signal differences across the TSS window, with higher values indicating greater confidence in differential enrichment between groups. **k)** Schematic of working model.

